# Networks of enhancers and microRNAs drive variation in cell states

**DOI:** 10.1101/668145

**Authors:** Meenakshi Chakraborty, Sofia Hu, Marco Del Giudice, Andrea De Martino, Carla Bosia, Phillip A. Sharp, Salil Garg

**Author notes:** = co-corresponding authors, and.

## Abstract

Cell-to-cell variation in gene expression is a common feature of developmental processes. Yet, it remains unclear whether molecular mediators can generate variation and how this process is coordinated across loci to allow the emergence of new cell states. Using embryonic stem cells (ESCs) as a model of development, we found interconverting cell states that resemble developmental expression programs and vary in activity at specific enhancers, such as those regulating pluripotency genes *Nanog* and *Sox2* but not *Pou5f1* (*Oct4*). Variable enhancers drive expression of variable genes, including those encoding microRNAs (miRNAs). Notably, variable miRNAs increase cell-to-cell variation by acting on neighborhoods of pluripotency genes. The encoded, variable pluripotency factors bind variable enhancers, forming a feedback loop that amplifies variation and allows the emergence of new cell states. These findings suggest gene regulatory networks composed of enhancers, protein-coding genes, and miRNAs harness inherent variation into developmental outcomes.

## Introduction

Pluripotent ESCs derived from the inner cell mass of the blastocyst contain the potential to form all cells of the adult vertebrate through sequential developmental stages. While external signals such as morphogens coordinate portions of this process, variation inherent to cells also plays a role, with small numbers of cells exhibiting spontaneous self-organization in the absence of external signals (Harrison et al., 2017; Rivron et al., 2018). Cell states are the expression of specific gene programs that often lead to a certain cell behavior. Variation in cell states is crucial for morphogenesis, as failure to transition between states leads to failures in development (Shahbazi et al., 2017). A prevailing view of cell-to-cell variation in gene expression attributes it to 'stochastic processes' at gene loci (Chang et al., 2008; Deng et al., 2014; Eldar and Elowitz, 2010; Kumar et al., 2014; Levine and Tjian, 2003; Raj and van Oudenaarden, 2008; Singh et al., 2010). However, an alternative hypothesis is that variation can be driven and coordinated across loci by gene regulatory elements and co-opted by the cell to enable state transitions. To evaluate this idea, we set out to measure heterogeneity in ESC states and define the molecular mediators that actively generate and coordinate variation in key pluripotency genes.

Pluripotency factors such as Pou5f1 (Oct4), Sox2, and Nanog (together ‘OSN’) are encoded by pluripotency genes and bind their own enhancers. High density clusters of binding sites have been termed super-enhancers (SEs), and OSN are known to bind together at SEs (OSN-SEs), forming a core transcriptional regulatory network that maintains ESC identity (Dowen et al., 2014; Hnisz et al., 2013; Kagey et al., 2010; Whyte et al., 2013). This regulatory network also contains microRNAs (miRNAs), small RNAs that bind and regulate genes in mammals through the effector protein Argonaute (Ago) (Marson et al., 2008). For example, the highly expressed miRNA cluster miR-290-295 maintains ESC identity during embryonic development (Medeiros et al., 2011; Melton et al., 2010). However, it is unknown whether this core regulatory network of enhancers, genes, and microRNAs plays a role in establishing the cell-to-cell variation necessary for subsequent development.

Here we studied naturally arising variation within ESCs by generating knock-in reporters at endogenous pluripotency genes. We found ESCs exhibit intrinsic heterogeneity through spontaneous formation of interconverting cell states that resemble distinct developmental expression programs. States vary in activity at particular enhancers that regulate variable, state-specific protein coding genes and miRNAs. Variable miRNAs such as miR-182 bind interaction neighborhoods of variable pluripotency genes such as *Nanog*, *Sox2*, and *Esrrb*. Notably, miRNAs amplify cell-to-cell variation in the expression of these genes, in contrast to previously described roles for miRNA in reducing gene expression variation. In turn, Nanog, Sox2, and Esrrb bind variable enhancers, suggesting these components act together with variable miRNAs in a network that amplifies inherent variation. In contrast, other well-expressed pluripotency genes such as *Pou5f1*, *Smad1*, and *Tcf3* are not bound by variable miRNAs, do not vary across ESC states, and do not show significant binding to variable enhancers. Together, our findings imply that the core transcriptional gene regulatory network of ESCs can be divided into at least two circuits, one that maintains ESC cell-type identity and one that coherently amplifies variation to achieve transition to a new cell state.

## Results

### ESC exhibit inherent variation that resembles developmental gene expression programs

To explore molecular mediators of ESC variation, we generated cells with heterozygous insertions of fluorophore tags at the endogenous loci of pluripotency genes *Nanog* and *Sox2*, joined by post-translational self-cleaving peptide sequences (*GFP*-P2A-*Nanog* and *Sox2*-T2A-*mCherry* respectively; see Fig. S1A). ESCs cultured in serum with leukemia inhibitory factor (LIF) showed remarkable heterogeneity in levels of Nanog and Sox2, clustering into three predominant states (Fig. 1A: State 1 = high Nanog and high Sox2, State 2 = low Nanog and high Sox2, State 3 = low Nanog and low Sox2), consistent with previous reports describing heterogeneity in Nanog levels across ESC populations (Chambers et al., 2007; Kumar et al., 2014; Singer et al., 2014). When cells from these distinct states were isolated by flow cytometric sorting and cultured identically (serum + LIF), each state recapitulated the heterogeneity of the parental population (Fig. 1A, bottom).

**Figure 1:**
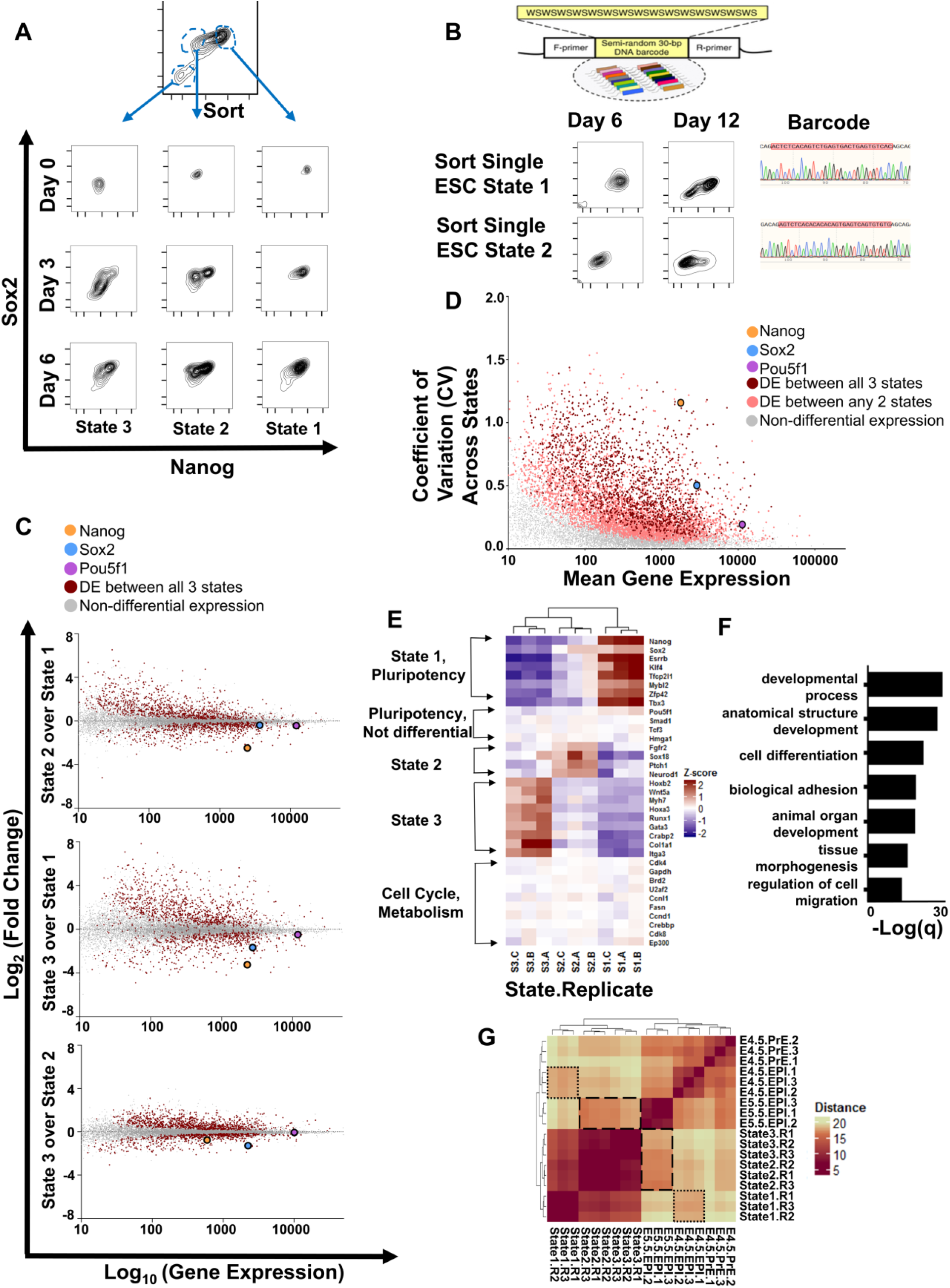
ESC fluctuate between cell states that vary in expression of developmental regulators. **A.** ESC labeled by heterozygous insertion of fluorophore tags at the endogenous loci for *Nanog* and *Sox2* (*GFP-Nanog*, *Sox2-mCherry*) were separated into three distinct cell states by flow cytometric sorting and cultured identically. The population is shown over time. **B.** A unique barcode was introduced into each ESC (Bhang et al., 2015). Single ESCs from States 1 and 2, respectively, were isolated and cultured. State distribution and sequencing of the barcode region (red highlight) are shown. **C.** Gene expression changes between ESC states for protein coding genes, shown as average of expected counts versus fold change between states (MA plots). Highlighted in red are genes with significantly differential expression between all three states. *Nanog*, *Sox2*, and *Pou5f1* (*Oct4*) are indicated. **D.** Coefficient of variation (CV)-mean plot of protein-coding gene expression across three states. Highlighted in red are genes with differential expression between all three states; in peach are genes with differential expression between any two states. *Nanog*, *Sox2*, and *Pou5f1* (*Oct4*) are indicated. **E.** Heatmap of normalized expression across ESC states for selected genes. Three biological replicates (A-C) are shown. **F.** Top gene ontology (GO) analysis terms for genes differentially expressed between all three states. FDR-q values for each ontology term are shown. See also Table 1. **G.** Heatmap of gene expression distance between ESC States 1-3 compared to expression profiles of embryonic development (E4.5 pre-epiblasts, E4.5 epiblasts, and E5.5 epiblasts, from (Boroviak et al., 2015)). Highlighted are State 1 vs E4.5 epiblasts (small dashes) and States 2-3 versus E5.5 epiblasts (large dashes).

**Table 1:** Selected gene ontology (GO) terms for processes enriched within ESC states. GO terms are shown for each state along with p values.

| State 1 | State 2 | State 3 |
| --- | --- | --- |
| Response to leukemia inhibitory factor ($p=1.99*10^{-7}$) | Epithelium development ($p=3.60*10^{-5}$) | Tube morphogenesis ($p=5.56*10^{-14}$) |
| Cellular response to cytokine stimulus ($p=1.55*10^{-4}$) | Ion transport ($p=2.10*10^{-5}$) | Animal organ development ($p=2.03*10^{-11}$) |
| Cell adhesion ($p=9.24*10^{-4}$) | Cell fate determination ($p=5.18*10^{-4}$) | Regulation of vasculature development ($p=3.64*10^{-14}$) |
| Cellular developmental process ($p=2.31*10^{-4}$) | Trans-synaptic signaling by neuropeptide ($p=9.67*10^{-4}$) | Regulation of angiogenesis ($p=2.18*10^{-13}$) |
| Stem cell population maintenance ($p=9.79*10^{-4}$) | Neuron recognition ($p=9.98*10^{-4}$) | Regulation of cartilage development ($p=9.59*10^{-9}$) |

Next, we assessed whether single ESCs also displayed an inherent capacity for heterogeneity by introducing a unique molecular barcode into individual ESCs, sorting single cells from a given state, and assessing their ability to repopulate the other states. Over time, single-cell-derived ESCs containing a unique barcode switched into other states (Fig. 1B). Analysis of an unsorted barcoded ESC population also indicated single cells were switching between states, as distinct cell barcodes observed only in one state on day 0 were observed in multiple states days later (Fig. S1B). Given these results, it is unlikely that sorted states recapitulate parental heterogeneity due to incomplete separation followed by differential growth rates. Rather, barcoding data indicate that state heterogeneity results from single cells switching states.

To gain insight into the observed ESC states, we characterized their coding and noncoding transcriptomes by ribosomal RNA-depleted RNA-sequencing (Ribo^−^ RNA-seq) (data: Supplemental Item 3). Protein-coding genes differentially expressed between all three states were highly enriched for developmental regulators and pluripotency genes (including *Nanog*, *Sox2*, and *Esrrb*) and depleted for housekeeping, cell cycle, and metabolic genes, suggesting variation across states was specific to developmental loci (Fig. 1C-1F and Fig. S2A-B). Notably, states displayed equally high expression of certain pluripotency genes such as *Pou5f1* (*Oct4*), *Smad1*, and *Tcf3* (Fig. 1C-1E), signifying that all three states are still embryonic stem cells but of differing functional predilection. In particular, ontology analysis of genes highest expressed within each state revealed that State 1 resembles a naïve, cytokine responsive population of cells, whereas State 2 displays increased expression of pre-ectodermal makers such as *Sox18* and *Neurod1* and State 3 displays increased expression of pre-endodermal and pre-mesodermal markers such as *Gata3* and *Hoxa3* (Table 1, Fig. 1E, Fig. S2C). To supplement the ontology analysis, we compared the highest expressed genes in each state to characterized gene expression profiles of the mouse blastocyst at developmental stages ranging from E4.5 to E5.5 (Boroviak et al., 2015; Shahbazi et al., 2017). We calculated the distance in gene expression between conditions (see Methods). While States 1-3 were most similar to each other, State 1 was closer in expression to E4.5 epiblasts than were States 2 and 3 whereas the latter were closer in expression to E5.0 or E5.5 (Fig. 1G and Fig. S2D, note increased red shading, meaning lower expression distance, at E4.5.EPI for State 1 replicates (small dashes) compared to States 2 and 3 replicates (large dashes) and vice versa for E5.5.EPI). Overall, we find that ESCs in culture contain inherent fluctuations observable as transitions between states that resemble developmental gene expression programs.

### A subset of super-enhancers varies between ESC states

We set out to determine the molecular nature of the variation between ESC states. Differences between cell types have been attributed to the activity of enhancers, specifically high-density super-enhancers (Whyte et al., 2013). We reasoned that differences between inter-converting cell states within a population may also be due to SE activity. Active enhancers are transcribed and produce transcripts known as enhancer RNAs (eRNAs). We define variable super-enhancers as those with differential activity in eRNA transcript production (da-SEs). We assessed eRNA production in States 1-3 across known enhancer regions marked by transcription factor binding and hallmarks of transcriptional regulation (Suzuki et al., 2017; Whyte et al., 2013). A remarkably high fraction of SEs defined by the chromatin mark H3K27Ac (H3K27Ac-SE) showed differential activity between ESC states, as did SEs defined through clusters of bound Pou5f1, Sox2, and Nanog (OSN-SE) (Fig. 2A). Moreover, in contrast to SEs that varied in activity between states, some strong SEs such as those driving *Pou5f1* were more consistently active across all three states (Fig. 2B and S3A, compare CV for *Nanog* & *Sox2* SEs to those for *Pou5f1* SE), indicating SE activity varies between cell states at a subset of core regulatory enhancers.

**Figure 2:**
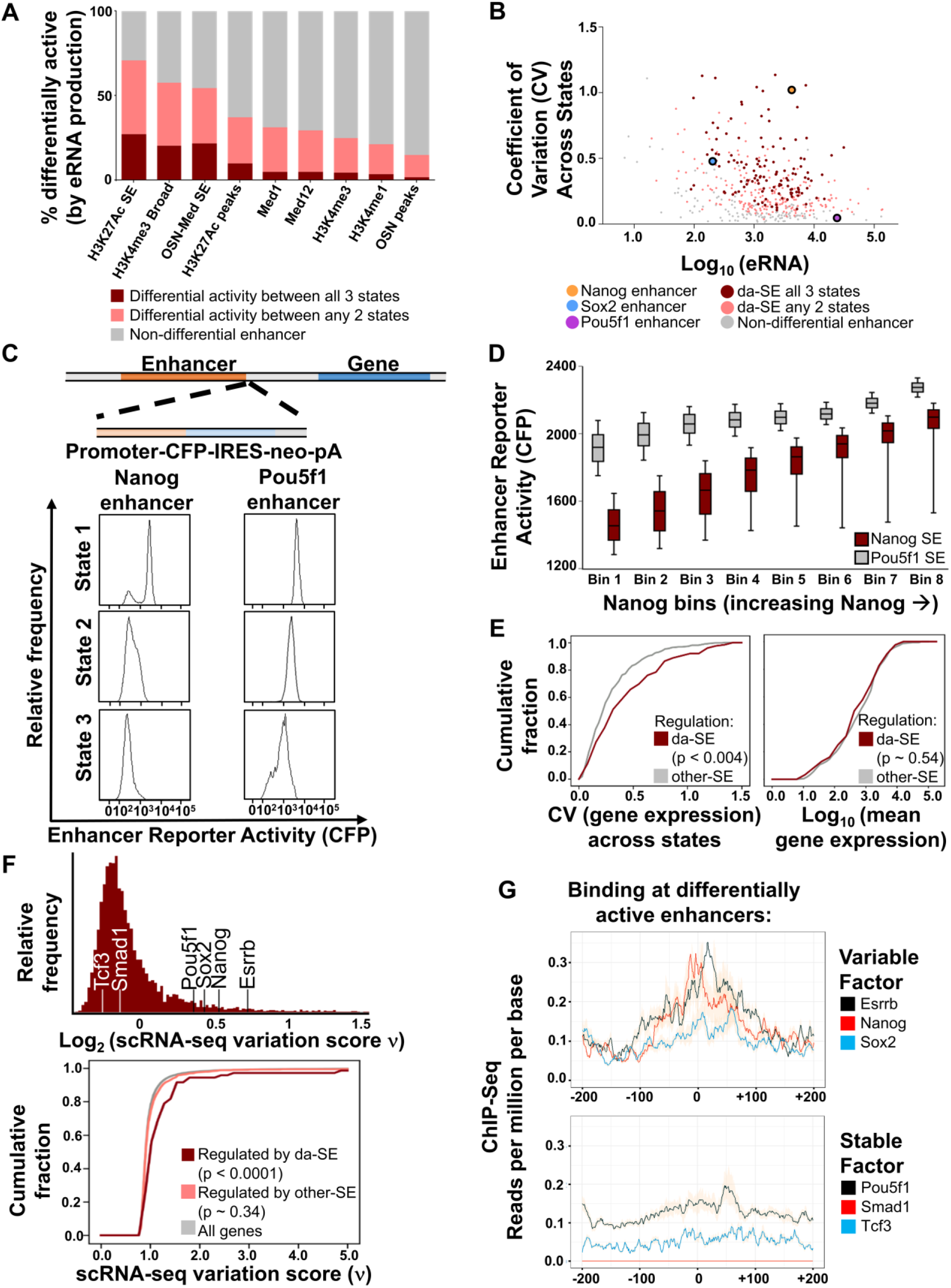
A subset of Super-Enhancers (SEs) varies in activity between ESC states. **A.** Proportion of each enhancer type showing differential activity between ESC states. **B.** CV-mean plot for SE activity across three states for H3K27Ac SEs. Highlighted in red are enhancers with significantly differential activity between all three states. In peach are SEs with differential activity between any two states; in gray are other SEs. Enhancers associated with *Nanog*, *Sox2* and *Pou5f1* are labelled. **C.** Reporters (promoter-*CFP*-IRES-*neo*-polyA) were inserted at enhancers controlling *Nanog* and *Pou5f1* in separate cell lines (which were also labelled at *Nanog* and *Sox2* loci, see Fig. S1A). Relative reporter activity (CFP) in States 1-3 is shown. **D.** Box-whisker plots for *Nanog* & *Pou5f1* enhancer activity (CFP fluorescence intensity) is plotted within the same narrow windows (“bins”) of Nanog expression ranging from Nanog-low to Nanog-high cell states. Whiskers extend from 25^th^ to 75^th^ percentile. **E. Left:** Cumulative Distribution Function (CDF) plot of gene expression variation across states (CV) for genes regulated by differentially active OSN-SEs (da-SE) and genes regulated by all other OSN-SEs (other-SE). **Right:** Average gene expression across three states for da-SE and other-SE regulated genes. **F. Top:**Distribution of single cell variation test statistic (ν) scores for 7,259 genes across 2,299 well-sampled cells measured by scRNA-seq. *Nanog*, *Sox2*, and *Esrrb* are indicated, as are *Pou5f1*, *Smad1*, and *Tcf3*. **Bottom:**CDF of single cell variation test statistic (ν score) for: all genes, da-SE regulated genes, and other-SE regulated genes. Kruskal-Wallis p-values are shown. See Methods for further details. **G.** ChIP-seq binding (reads per million mapped per base) at da-SEs. Each enhancer was extended to a minimum of 1 kb, all enhancers were scaled to 1000 bins, and reads normalized to input. Shown are the middle 400 bins of binding for the indicated genes. **Top**: Variable factor binding at da-SE and **bottom**: Less variable (stable) factor binding at da-SE.

To directly measure the variability in SE activity between states, we inserted a fluorescent reporter (Cerulean, or CFP) at SEs and measured both enhancer activity and cell state by flow cytometry, similar to a previously reported strategy (Stelzer et al., 2015). We report SEs where insertion of the reporter did not significantly alter cell states. We found high state-to-state variation of a SE controlling the variable pluripotency gene *Nanog* (*Nanog*-SE) (Fig. 2C). In contrast, activity at an SE controlling the non-variable pluripotency gene *Pou5f1* (*Pou5f1*-SE) was almost identical between State 1 and State2 (Fig. 2C), though this enhancer did show some reduced activity in State 3. Even within a single state, *Nanog*-SE variation was greater than *Pou5f1*-SE variation (Fig. 2C, note the bimodal peak for *Nanog*-SE activity in State 1 compared to one peak for *Pou5f1*). Further, we analyzed variation within state by plotting reporter activity within narrow windows (bins) of Nanog expression. We found *Nanog*-SE activity varied more than *Pou5f1*-SE activity when compared in this fashion (Fig. 2D, note the taller box and whisker plots for *Nanog*-SE compared to *Pou5f1*-SE indicating higher variance or 'spread' of the data). Similarly, we inserted reporters separately at the SEs controlling the variable pluripotency gene *Esrrb* and the non-variable pluripotency gene *FGF4*. Analogously to the result for *Nanog* and *Pou5f1*, we found relatively higher variation for *Esrrb*-SE activity compared to *FGF4*-SE activity (Fig. S3B). Together, these data demonstrate that enhancers exist in a ‘variation hierarchy’ whereby a subset are inherently more prone to variation in activity between and within cell states.

### Transmission of enhancer variation to regulated genes

As enhancers are core regulatory components for gene expression, we reasoned that state-to-state variation in activity at SEs could in part account for the variability in genes that were differentially expressed between states. To assess this possibility, we first tried deleting or interfering with SEs using CRISPR-Cas9 targeting but found this either resulted in ESC differentiation or grossly changed the gene expression distributions of States 1-3 (data not shown), complicating assessment of the subtle fluctuations between states under consideration in our study. Instead, we utilized maps defining the regulated genes of OSN-SEs (Dowen et al., 2014). First, we separated OSN-SEs into those differentially active between all three states and compared them to all other SEs (da-SEs and other-SEs, respectively, note that these OSN-SE categories are used throughout the remainder of the study). Next, we plotted the coefficient of variation (CV) across states for the regulated genes. We found a significant shift towards higher CV for genes regulated by da-SEs compared to other-SEs (Fig. 2E, left). Importantly, this shift was not due to different expression levels for genes regulated by da-SEs (Fig. 2E, right). Further, da-SE were most active in the same state where their regulated genes were most highly expressed (Fig. S3C-D). This supported the notion that variation between states for differentially expressed genes arose in part from enhancers with variable activity between states.

In addition to variation between states, we observed variation within state for da-SE such as the *Nanog*-SE (Fig. 2C-D). Both between-state and within-state heterogeneity require single cells to vary from one another. Additionally, measurements of gene expression across cells in bulk populations can obscure effects on single cells missed by taking ensemble averages. Therefore, we performed single cell RNA-sequencing (scRNA-seq) to relate variation at enhancers to gene expression variation across single cells. We identified highly variable genes using a previously developed test statistic (ν) that corrects the coefficient of variation for technical sampling noise present in scRNA-seq experiments (Klein et al., 2015). We found highly variable genes included da-SE-regulated *Nanog*, *Sox2*, and *Esrrb* (Fig. 2F, top). In contrast, *Pou5f1* showed a moderate degree of variation and both *Smad1* and *Tcf3* were less variable across single cells (Fig. 2F, top). Indeed, genes regulated by da-SEs showed significantly higher variation across single cells than genes regulated by other-SEs (Fig. 2F, bottom). Together these results suggest that da-SE can drive cell-to-cell variation in their regulated genes.

### Feedback of variable pluripotency genes on variable enhancers

Variation in molecular components of a gene regulatory network could either add together or cancel out, similar to constructive or destructive interference of waves. Given that heterogeneous states reform each other when separated (see Fig. 1), we reasoned fluctuations at variable molecular components in ESCs may be wired together, allowing them to exhibit constructive properties in subsets of cells. The core transcriptional regulatory network of ESCs is known to contain binding of pluripotency factors such as OSN at enhancers; however, whether binding can be functionally subdivided on the basis of cell-to-cell variation has not been explored. Hence, we assessed the binding of variable pluripotency factors Nanog, Sox2, and Esrrb and less variable pluripotency factors Pou5f1, Smad1, and Tcf3 at both da-SEs and other-SEs. We found a significant degree of binding of Nanog, Sox2, and Esrrb to da-SE (Fig. 2G, top), whereas Pou5f1 and Tcf3 did not show a distinct binding peak to da-SE and Smad1 did not bind enhancers at all (Fig. 2G, bottom and Fig. S3E). Variable pluripotency factors and less variable pluripotency factors Pou5f1 and Tcf3 showed similar binding to other-SEs (Fig. S3E). This is consistent with a model whereby in a subset of ESCs variable pluripotency factors such as Nanog bind and guide modification of variable enhancers, which then feed forward on variable genes, priming this subset of cells for transition to a new state. In the meantime, all pluripotency factors can function together at relatively stable SEs to maintain ESC cell type identity. Thus, within the ESC core regulatory network there may be a sub-network of elements forming a circuit capable of amplifying inherent variation.

### The variable ESC network contains miRNAs

In addition to driving the expression of protein-coding genes, enhancers are known to control the expression of miRNAs. miRNAs are intriguing candidate cell state controllers because individual miRNAs can regulate hundreds of genes, which could allow cell-to-cell fluctuations in miRNA to generate relatively large effects on cell state (Garg and Sharp, 2016). To examine this possibility, we first determined miRNA expression in each state by small RNA-sequencing (data: Supplemental Item 4) and noted differentially expressed miRNAs (DE-miRNA) (Fig. 3A-B). DE-miRNA did not include most members of the miR-290-295 family, which were more equally expressed in all three states. Instead, DE-miRNA represented many miRNAs with no previously characterized function in ESC. We found that DE-miRNA were enriched for regulation by da-SE (Fig. 3C), consistent with the idea that this group of miRNA is part of a variable genetic circuit.

**Figure 3:**
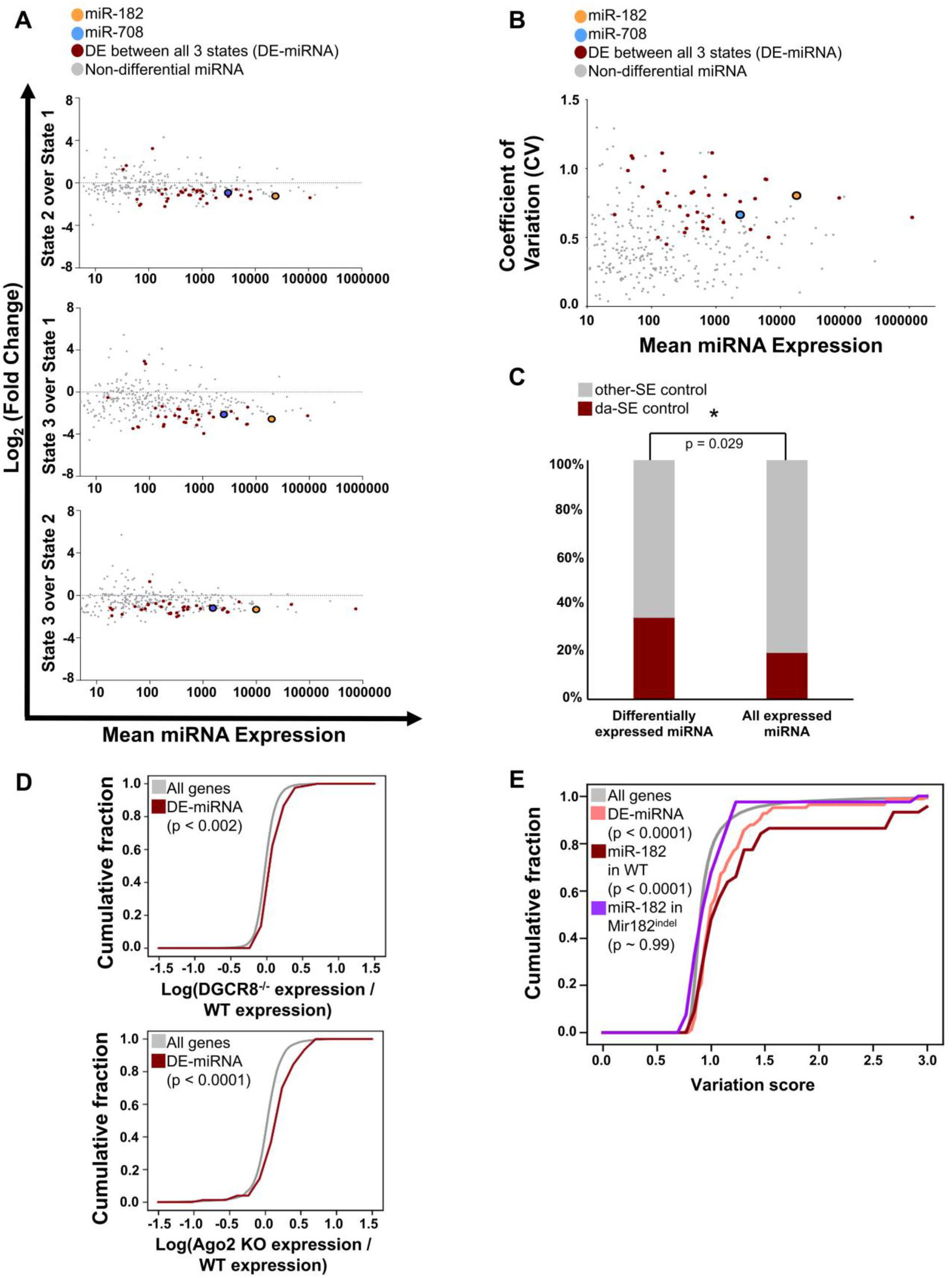
Variable microRNAs increase cell-to-cell variation of their target genes. **A.** Changes between ESC states for miRNAs, determined by small-RNA sequencing of sorted States 1-3. Expected counts versus fold change between states is plotted (MA plot). Highlighted in red are miRNAs with differential expression between all three states (DE-miRNA). miR-182 and miR-708 are indicated. **B.** CV-mean plot for miRNA expression across states. DE-miRNA are marked in red. miR-182 and miR-708 are indicated. **C.** Proportion of miRNA controlled by differentially active vs. other enhancers for both DE-miRNA and all other miRNAs. miRNA from genes not under enhancer control are omitted from analysis. Hypergeometric p-value for enrichment is shown. **D. Top:** CDF for the ratio of gene expression in DGCR8^−/−^ ESC vs WT ESC for all genes or genes targeted by ≥ 2 DE-miRNAs. **Bottom**: CDF for the ratio of gene expression in Ago2-inducible ESC competent for miRNA activity (1 μg/mL doxycyline) versus no miRNA activity (0 μg/mL doxycycline for 48 hours). Expression for all genes was measured by bulk RNA-seq. Kolmogorov-Smirnov (K-S) p-value is shown. **E.** CDF of variation score across single cells (ν score) for all genes (gray), genes targeted by ≥2 DE-miRNAs (peach), genes targeted by miR-182 in WT ESC (red), and genes targeted by miR-182 in *Mir182*^*indel*^ ESC (violet). Kruskal-Wallis p-values are shown.

To gain insight into the function of DE-miRNAs, we analyzed Argonaute-miRNA cross-linking and immunoprecipitation (CLIP) to targets (Bosson et al., 2014). In these cells, FLAG-HA-hAGO2 expression is controlled from a doxycycline inducible transgene in an endogenous Ago1^−/−^/2^−/−^/3^−/−^/4^−/−^ knockout background (Ago2-inducible ESC), and in the absence of Ago2 induction miRNA activity is lost (Zamudio et al., 2014). This allows high resolution definition of Ago2-miRNA complexes and their bound targets by analyzing clusters of Ago2 binding. We assigned each cluster to a particular miRNA by considering a combination of site type affinity and miRNA expression (see Methods). We found transcripts bound by DE-miRNA showed a higher number of Ago2-miRNA binding events per each mRNA target (Fig. S4A). Moreover, binding was skewed toward lower affinity ‘6-mer’ miRNA site type matches and away from higher affinity ‘8-mer’ sites (Fig. S4A). Interestingly, we found a high degree of DE-miRNA binding to *Nanog*, *Sox2*, and *Esrrb* transcripts (Fig. S4B), pluripotency genes not previously appreciated to be under miRNA regulation. We did not find evidence of significant binding of DE-miRNA to less variable pluripotency gene *Pou5f1*, *Smad1*, or *Tcf3* transcripts.

### Variable miRNAs increase variation of target mRNAs

To assess whether loss of DE-miRNA impacts their mRNA targets, we analyzed ESC where the microRNA biogenesis and effector machinery is disrupted. For example, DGCR8 controls miRNA processing from precursor ‘pri-miRNA’ to ‘pre-miRNA’ forms (Gregory et al., 2004; Wang et al., 2007). We found modestly increased mean mRNA levels (‘de-repression’) for targets of multiple DE-miRNAs (≥2 clusters of Ago2 binding) in DGCR8^−/−^ ESC (DGCR8 KO) (Fig. 3D, top). Next, we analyzed DE-miRNA targets in Ago2-inducible ESC after withdrawal of doxycycline. Consistent with results for DGCR8^−/−^, DE-miRNA targets showed modest de-repression in Argonaute deficient ESC (Fig. 3D, bottom), indicating DE-miRNA can have a suppressive effect on their targets. We sought to extend these results to individual DE-miRNA. We generated ESCs deficient in either DE-miRNA miR-182 or DE-miRNA miR-708 by inducing CRISPR-Cas9 targeted indels in the hairpin loop of the miRNA gene (*Mir*^*indel*^) (Chen et al., 2015). We confirmed *Mir*^*indel*^ ESCs represented miRNA knockouts using measurement of miRNA levels and functional reporter assays (Fig. S4C) (Mukherji et al., 2011). Surprisingly, there was no difference in average expression levels across all cells for miR-182 and miR-708 bound transcripts in *Mir182*^*indel*^ and *Mir708*^*indel*^ ESCs respectively when compared to WT ESC (Fig. S4D). However, it is possible a regulatory effect was masked due to considering average expression across all cells, leaving open the possibility these miRNAs were regulating targets in a subset of cells.

Intriguingly, we noted DE-miRNA bound genes, including miR-182 bound genes, showed high variation across single cells in WT ESC (Fig. 3E). To determine whether miR-182 impacted variation of its targets, we performed scRNA-seq on *Mir182*^*indel*^ ESC in parallel to WT ESC. We compared variation of miR-182 bound targets in these two cell types by ν score. Strikingly, loss of miR-182 resulted in a sharp reduction in variation of miR-182 bound genes across single cells (Fig. 3E, compare left shift of miR-182 bound targets in *Mir182*^*indel*^ ESC (violet) to miR-182 targets in WT ESC (maroon)) without resulting in significant changes in gene expression across single cells (Fig. S4E). Thus, while miR-182 did not have a detectable effect on average target expression across all cells, it appeared to have a significant effect on cell-to-cell variation of its targets. Previous studies have highlighted roles for miRNA in reducing gene expression variation of their targets compared to genes without miRNA regulation. However, cell-to-cell variation in the miRNA pool could result in transmission of this variation to bound targets (Garg and Sharp, 2016; Schmiedel et al., 2015). In the context of a gene regulatory network our findings imply miR-182 could work in this fashion in ESC, serving to increase cell-to-cell variation at its bound targets in a subset of cells.

### Some pluripotency gene neighborhoods are highly bound by variable miRNAs

Cell states are defined by coordinated expression of gene groups, meaning that transitions between states require coordinated variation of genes. We sought to gain insight into whether variation at highly variable genes was coordinated and related to the observed states. To this end, we used network inference methods to construct gene interaction neighborhoods from our scRNA-seq data. We chose a method based on mutual information, whereby the neighborhood of a chosen ‘node’ gene represents the set of genes most closely correlated with it and with each other, similar to a previous analysis in ESC (Klein et al., 2015; Li and Horvath, 2007) (see also Methods). The emphasis of this method on topology helps alleviate artifacts in correlation strength that may arise from the low technical sampling of transcripts in scRNA-seq. The neighborhoods of *Nanog*, *Sox2*, and *Esrrb* are shown (Fig. 4A and Fig. S5A). We confirmed these *in silico* inferred neighborhoods were meaningful by testing the covariation of Nanog with the encoded proteins of neighbors *Eif2s2*, *Esrrb*, and *Hsp90ab1*. For each of these genes, we introduced an additional fluorophore tag at its endogenous locus in *Nanog* fluorophore-tagged cells. Eif2s2, Esrrb, and Hsp90ab1 all showed covariation with Nanog by this method (Fig. S5B). Further, the neighborhoods of *Nanog*, *Sox2*, and *Esrrb* all contain each other as members and have additional mutual neighbors (Fig. S5C), supporting the idea that these genes interact and form an interconnected clique.

**Figure 4:**
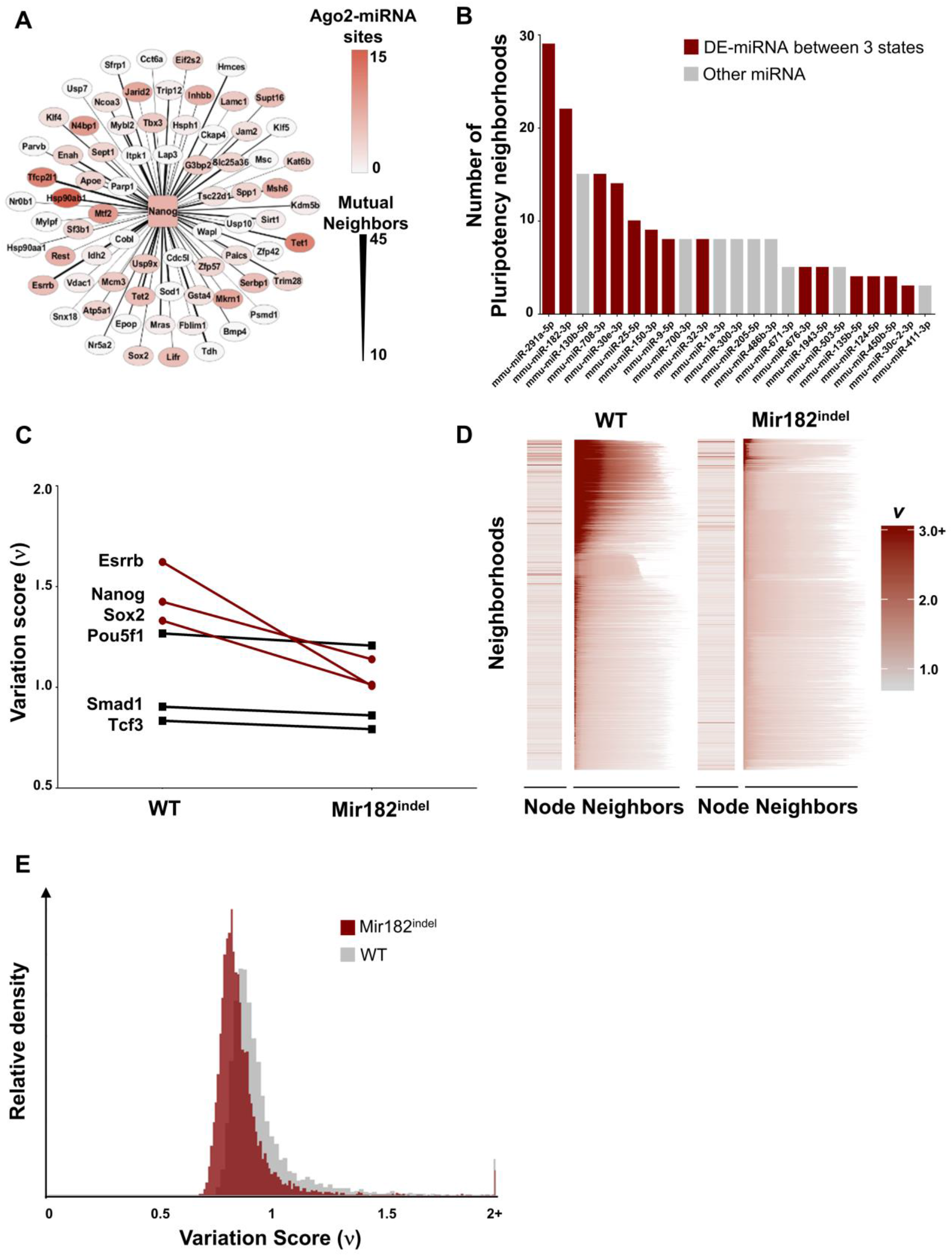
Coordination of variation across gene neighborhoods bound by variable miRNAs. **A.** *Nanog* interaction neighborhood, constructed by analyzing covariation across single cells (see Methods). Thickness of *Nanog* connection represents number of mutual neighbors between *Nanog* and that gene, red shading indicates degree of Argonaute binding (miRNA activity). **B.**Number of pluripotency neighborhoods (out of 55) significantly enriched for binding by each miRNA. DE-miRNAs are marked in red. **C.** Variation score across single cells (ν) for variable pluripotency genes *Nanog*, *Sox2*, and *Esrrb* and less variable (stable) pluripotency genes *Pou5f1*, *Smad1*, and *Tcf3* in WT vs. *Mir182*^*indel*^ ESCs. **D.** Variation scores (ν) for all non-empty neighborhoods in WT and *Mir182*^*indel*^ ESC. The variation score of the ‘node’ gene is shown at left, followed by the scores for all neighbors in the final neighborhood (arranged from highest variation score at left to lowest at right). Neighborhoods are plotted top to bottom by decreasing average variation of all neighbors for WT and *Mir182*^*indel*^ ESC separately. **E.** Overlaid histogram of variation (ν scores in WT cells and *Mir182*^*indel*^ cells for 6,107 genes well-sampled in both WT and *Mir182*^*indel*^ cells.

Next, we assessed the *Nanog* network for miRNA activity. We found remarkably high binding of the *Nanog* neighborhood by miRNAs (Fig. 4A). Similarly high miRNA binding was noted for the neighborhoods of other variable pluripotency genes, including *Sox2* and *Esrrb* (Fig. S5D). Further, we calculated whether neighborhoods were enriched for binding by particular miRNAs by comparing them to matched control neighborhoods. Control neighborhoods were constructed to contain the same number of genes of similar expression distribution and total miRNA binding as the neighborhood under interrogation (see Methods). Several miRNAs were enriched for binding within neighborhoods of variable pluripotency genes by this method (Fig. 4B). This group of miRNAs included many DE-miRNAs, including miR-182 and miR-708. Notably, neighborhoods of less variable pluripotency genes such as *Pou5f1*, *Smad1*, and *Tcf3* did not show high binding by miRNAs and contained fewer mutual neighbors than variable pluripotency genes (Fig. S5C, S5E). Partially distinct sets of miRNAs were enriched for binding other ontologically defined groups of interaction neighborhoods, such as cytoskeleton genes (Fig. S5F). The consistent enrichment of particular miRNAs within neighborhood groups defined by ontology suggests a role for miRNA in regulating these neighborhoods.

### Coordination and propagation of variation across neighborhoods

Coordinate regulation of gene neighborhoods by miRNAs could provide a mechanism for individual miRNAs to impact the variation of many genes. Namely, miRNAs could impact the variation of genes that they do not bind directly by binding and regulating their interacting neighbors. First, we compared the variation across single cells for DE-miRNA-bound genes *Nanog*, *Sox2*, and *Esrrb* in WT vs. *Mir182*^*indel*^ ESC. Indeed, DE-miRNA-bound genes had lower ν scores in *Mir182*^*indel*^ ESC than in WT, even if they were not direct targets of miR-182. By contrast, ν scores for non-DE-miRNA-bound genes *Pou5f1*, *Smad1*, and *Tcf3* were unchanged in *Mir182*^*indel*^ ESC vs. WT (Fig. 4C). This suggested that miR-182 propagates variability through the neighborhoods it binds in addition to promoting variability of its direct targets. To further assess whether variation can be propagated across neighborhoods, we plotted ν scores of all genes and their neighbors (Fig. 4D). In the leftmost column, we show the variation of the 'node' gene. Next, in each row we plot the ν score of each node gene's neighbors decreasing left to right, noting that neighborhoods differ in size (see Methods). We list genes in order of decreasing average ν score across their neighborhoods. Remarkably, highly variable genes show a strong tendency to group into the same interaction neighborhoods (Fig. 4D, note that red shading indicating higher ν score is clustered towards the top of the graph in a subset of neighborhoods rather than being evenly distributed across all neighborhoods). Overall, highly variable genes show synchronous co-variation when measured across a population of cultured cells. This indicates that individual genes do not vary stochastically with respect to each other. Rather, variation is organized at the level of neighborhoods. The degree and concentration of variation is significantly decreased in *Mir182*^*indel*^ ESC (Fig. 4D-E), consistent with the idea that miR-182 loss reduces cell-to-cell variation of bound targets, which in turn leads to less variation across *Mir182*^*indel*^ ESC neighborhoods compared to WT ESC neighborhoods.

### Variation in miRNA can drive variation in ESC states

Fluctuation in DE-miRNA levels causing fluctuations at highly variable genes in key neighborhoods could lead to a state transition and therefore an inherent propensity to reform a heterogeneous distribution of cells, such as observed in our study (see Fig. 1). This raised the possibility of miRNA regulation of *Nanog* and *Sox2* contributing to the observed distribution of cell states through variation of miRNA levels. First, we explored this possibility by constructing mathematical models involving *Nanog*, *Sox2*, and miRNA that could generate the observed distribution of cell states. We found that a minimal model in which *Nanog* and *Sox2* were regulated by two distinct miRNA pools could recapitulate the observed distribution of cell states solely through addition of cell-to-cell fluctuations in miRNA levels (Fig. 5A and Theory Note in Methods). The model predicted loss of variable miRNAs would cause loss of variation in cell states (Fig. 5A).

**Figure 5:**
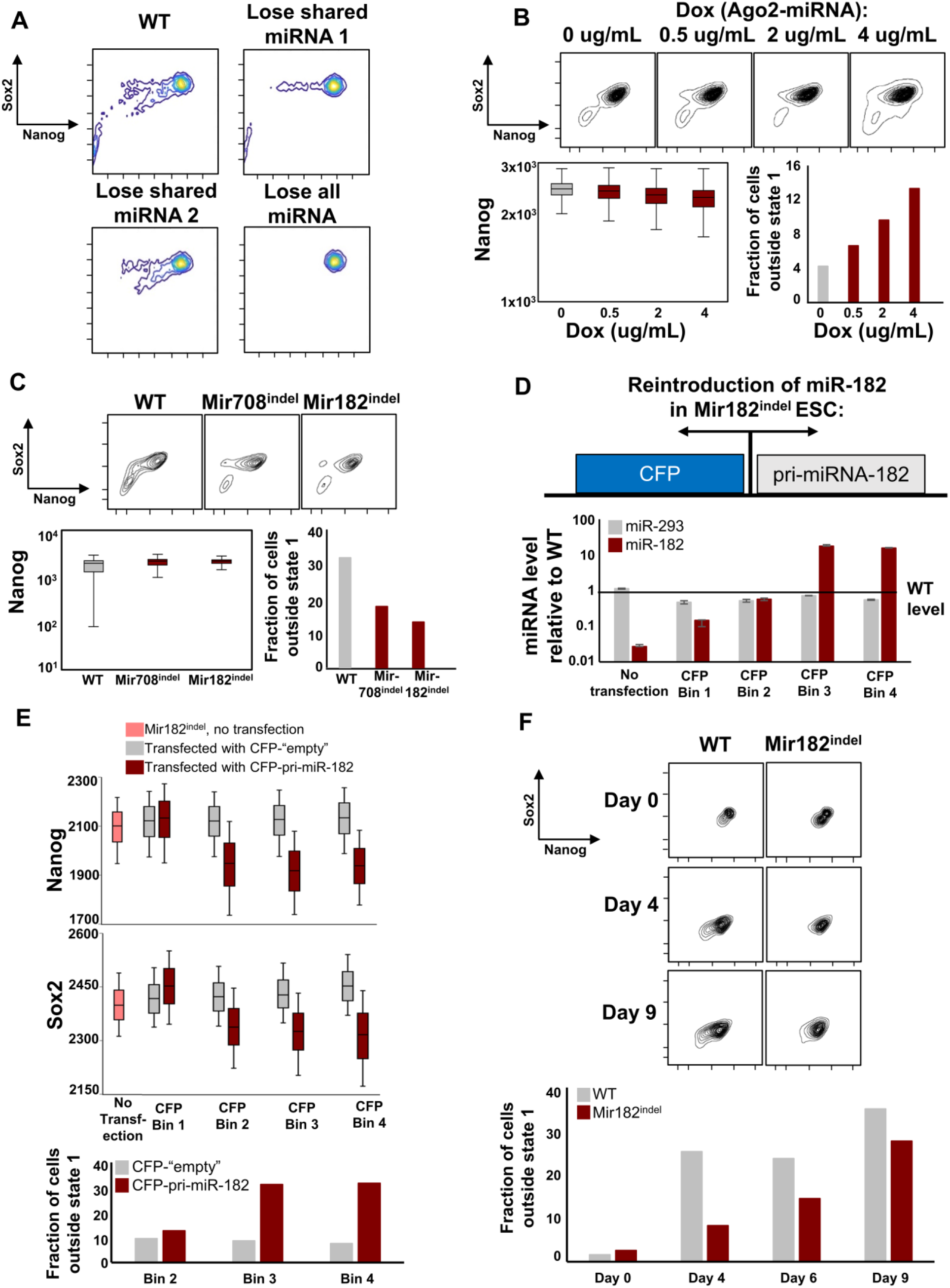
Variation in microRNA can drive variation in ESC states. **A.** Theoretical cell state distributions predicted by a mathematical model involving *Nanog*, *Sox2*, and miRNA in which cell-to-cell variation is added only in miRNA levels (see Theory Note). “Shared miRNA” refers to a miRNA that regulates both *Nanog* & *Sox2*. **B.** Cell state distributions in Ago2-inducible ESC (see Fig. 3D above) labelled at *Nanog* and *Sox2* loci by fluorophores (Fig. S1). Cells are cultured for 48 hours in the indicated concentrations of doxycycline (Ago2-miRNA). Variation in Nanog and the fraction of cells outside of State 1 are shown. See also Fig. S6A. **C.** Cell state distributions in WT, *Mir182*^*indel*^ and *Mir708*^*indel*^ ESC labelled at *Nanog* and *Sox2* loci by fluorophores (Fig. S1). Variation of Nanog and the fraction of cells outside of State 1 are shown. See also Fig. S6B. **D.** Reintroduction of miR-182 into *Mir182*^*indel*^ cells (from Fig. 5C above). *Mir182*^*indel*^ cells were transfected with a bi-directional expression plasmid expressing pri-miR-182 tightly coupled to Cerulean Fluorescent Protein (CFP) or with a plasmid containing CFP alone (‘empty’ control). Cells were isolated by flow cytometric sorting in increasing ‘bins’ of CFP expression and the levels of miR-182 and miR-293 quantified and plotted relative to WT ESC. **E.** Box-and-whisker plots for Nanog & Sox2 levels (GFP and mCherry fluorescence) in *Mir182*^*indel*^ cells plotted by CFP bins (miR-182 levels) as in part **D**. Note miR-182 levels do not change in ‘empty’ transfected cells (data not shown). See also Fig. S6C. **F.** State 1 WT and *Mir182*^*indel*^ ESCs isolated by flow cytometric sorting and cultured. The fraction of cells outside State 1 is shown. See also Fig. S6D.

To test this idea, first we inserted fluorophore tags at *Nanog* and *Sox2* (Fig. S1) into our Ago2-inducible cells (see Fig. 3D above) and withdrew doxycycline. With no induction of Ago2 and therefore no miRNA activity (doxycycline 0 μg/mL), cells exhibited relatively little variation in Nanog/Sox2 cell states and consequently relatively little variation in Nanog levels (Fig. 5B), in agreement with our model. As Ago2-miRNA levels were increased in these cells (doxycycline 0.5 - 4 μg/mL), we observed a titratable increase in cell state variation as a higher fraction of cells exited State 1, the dominant state (Fig. 5B), and transitioned to States 2-3 or became intermediate between states (Fig. S6A). These transitions also led to a measurable increase in Nanog variation across the population (Fig. 5B). Next, we tested whether loss of individual DE-miRNA could impact cell state variation. We inserted fluorophore tags at *Nanog* and *Sox2* into *Mir182*^*indel*^ and *Mir708*^*indel*^ ESC and assessed their cell state distributions. Strikingly, loss of individual DE-miRNA reduced cell state variation, with a measured reduction in the variation of Nanog (Fig. 5C, note shorter box and whisker plots for *Mir*^*indel*^ ESC compared to WT ESC) and reduction in cells outside of State 1 (Fig. 5C and Fig. S6B). This was consistent with the predictions of our model, and together these results established that loss of miRNA can reduce cell state variation in ESC.

We sought to test whether we could restore variation in *Mir*^*indel*^ cells by re-introducing the lost DE-miRNA. First, we constructed an inducible bidirectional expression plasmid expressing miR-182. Specifically, the plasmid expresses pri-miR-182 tightly coupled to a cyan fluorescent protein derivative (CFP), allowing close approximation of miRNA levels through levels of the fluorophore reporter. We transfected *Mir182*^*indel*^ cells containing reporters at *Nanog* and *Sox2* loci with this construct, allowing single cell measurement of miR-182, Nanog, and Sox2 levels through their respective fluorophores (CFP, GFP, mCherry, respectively). CFP levels tracked miR-182 expression and the latter was restored to approximately wildtype levels (in ‘Bin 2’) or overexpressed (in ‘Bins 3-4’, Fig. 5D). For comparison, we measured miR-293 in these bins and found it relatively unchanged. As an additional control, we transfected *Mir182*^*indel*^ cells with a *CFP*-only ‘empty’ plasmid in parallel to re-expression of miR-182. Compared to ‘empty’ control, *Mir182*^*indel*^ cells with miR-182 re-expressed showed an increase in cells outside State 1 (Fig. 5E and Fig. S6C), confirming restoration of miR-182 could partially restore cell state variation. Additionally, when comparing across or within bins, *Mir182*^*indel*^ cells in which miR-182 was restored showed increased variation in Nanog and Sox2 levels compared to control cells (Fig. 5E, note that mean levels of Nanog and Sox2 change across red ‘miR-182’ bins but stay similar across gray ‘empty’ bins).

Notably, this system involved adding variation to miRNAs outside the context of the variable genetic circuit. This is because plasmid transfection and induction efficiency are not uniform across the population, leading to differences in restored miR-182 levels between *Mir182*^*indel*^ cells transfected with pri-miR-182 expression plasmid. Thus, our results indicate that adding exogenous variation in miRNA levels can increase variation in cell states, and in Nanog and Sox2 levels. To further test the idea that miR-182 facilitated cell state transitions, we isolated State 1 ESC from WT and *Mir182*^*indel*^ ESC by flow cytometric sorting and assessed their ability to repopulate States 2-3 and intermediate states over time. In sorted cells, we found a delay in *Mir182*^*indel*^ State 1 exit compared to WT (Fig. 5F and S6D). Together, these findings supported the idea that within the variable genetic circuit fluctuations in miR-182 across cells led to miR-182 facilitated transition between states for a subset of cells. This could account for the increased variation in Nanog and Sox2 in WT ESC compared to *Mir182*^*indel*^ ESC as well as the increased variation across single cells for miR-182 targets (Fig. 3E). As this subset of cells exiting State 1 is a small portion of the overall ESC population at any given point in time this might also account for the lack of repression observed for direct miR-182 targets in bulk RNA-sequencing (Fig. S4D). We conclude that cell-to-cell fluctuations in DE-miRNA levels can directly contribute to variation in their bound pluripotency genes and interacting neighbors. In the context of a genetic circuit for variation comprising enhancers, miRNAs, and pluripotency genes, miRNAs can either transmit upstream variation to pluripotency genes or might themselves constitute a source of cell-to-cell variation. The coordination of variation at these elements in a subset of cells could enable transitions to new states and lead to a propensity for heterogeneity as a result of the variable gene regulatory network co-opting inherent fluctuations.

### Nanog and Sox2 capture a large portion of ESC state diversity

In principle, heterogeneity in ESC states could be defined by many different combinations of variably expressed genes. We defined cell states in ESC through their relative expression of *Nanog* and *Sox2*. To determine what proportion of heterogeneity in ESC states was captured by *Nanog* and *Sox2* reporters, we sought to represent ESC states in a gene unbiased manner by dimensionality reduction of our scRNA-seq data and to compare this representation to that given by States 1-3. We sought a method that would represent both local and global data structure in addition to emphasizing progression trajectories in ESC state space. Thus, we utilized PHATE (Potential of Heat diffusion for Affinity-based Trajectory Embedding) dimensionality reduction (Moon et al., 2019). We applied PHATE to the combined scRNA-seq data from WT and *Mir182*^*indel*^ ESCs while adding corrections for batch effects and depth of sampling per cell.

We plotted the first two PHATE coordinates while pseudocoloring WT and *Mir182*^*indel*^ cells by their enrichment for State 1-3 expression programs (Fig. 6, note WT cells are plotted white to blue and *Mir182*^*indel*^ cells plotted white to red by their enrichment for States 1-3, see Methods). We noted that State 1-3 programs, defined by expression levels of Nanog and Sox2, capture the majority of variation between cells along the PHATE axes (Fig. 6, note the progression of enrichment for the State 1 program at bottom left to the State 3 program at top right). Qualitatively similar results were obtained for dimensionality reduction by principle component analysis and a force directed graph based method (Fig. S7A-B, based on (Paul et al., 2015)). Importantly, we did not find significant clustering of cells when plotted by cell cycle, confirming that the observed heterogeneity was not due simply to variations in cell cycle (Fig. S7C). Since PHATE representations are made without choosing any particular gene as a starting point, we infer ESC cell states defined by Nanog and Sox2 capture a significant amount of the heterogeneity present.

**Figure 6:**
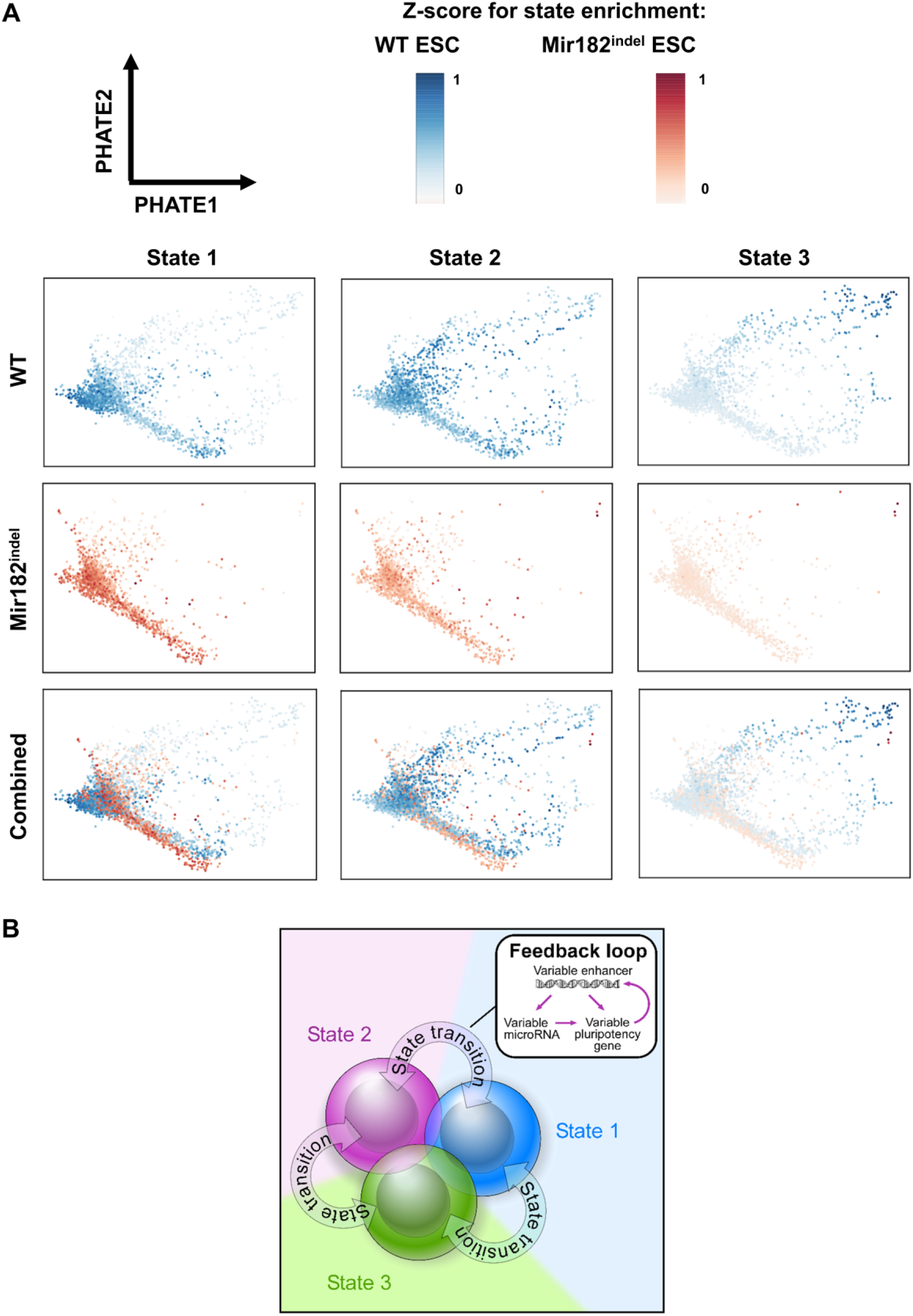
Nanog and Sox2 capture a large portion of ESC state diversity. **A.** PHATE dimensionality reduction applied to scRNA-seq data from both WT and *Mir182*^*indel*^ ESC together (see Methods). Cells are plotted according to their low dimensional representations (PHATE1 vs PHATE2). Each cell is colored for relative enrichment by gene expression signatures of States 1-3 for WT ESC (blue, top row) and *Mir182*^*indel*^ ESC (red, middle row) or both ESC types together (bottom row). **B.** Model for variation feedback loop in ESC leading to state transitions.

## Discussion

We find that ESCs exhibit intrinsic heterogeneity by spontaneously forming interconverting cell states that resemble developmental expression programs. States display distinct activity of enhancers along with distinct expression of protein-coding genes and miRNAs. Variable enhancers, pluripotency genes, and miRNAs all form an interconnected variable genetic circuit (Fig. 6B), suggesting a mechanism by which variation can amplify across a gene regulatory network and result in a new cell state.

Together our results provide a framework for how cell states can emerge from the organization of inherent fluctuations by a gene regulatory network but leave open the question of where and how fluctuations arise in the first place. Gene expression is known to occur in discrete bursts of transcription, and enhancers have been connected to the modulation of bursting for their regulated genes (Fukaya et al., 2016; Larsson et al., 2019). Additionally, enhancers have been proposed to form phase condensates (Cho et al., 2018; Chong et al., 2018; Hnisz et al., 2017; Sabari et al., 2018), and these may contribute fluctuations. Future work further defining the structural and regulatory features of different enhancer classes that vary in activity between loci will be of great interest. We found variable miRNAs were more likely to bind targets weakly and multiply, a hallmark of interactions more susceptible to fluctuation. Perhaps miRNA variation will also prove a key source of fluctuation for cells to co-opt in state transitions. Future studies focused on the mechanisms by which “microscale inhomogeneities” (Chen et al., 2018) can form cell-to-cell will be necessary to fully understand the fluctuations within ESC.

While the present study focuses on enhancers and miRNAs, many other molecular mediators are likely to contribute to cell-to-cell variation. Of particular interest may be RNA binding proteins or splicing factors, as these regulators can also impact the expression of many genes, enabling state transitions from fluctuations in a few key mediators. Our results suggest a subset of the core ESC gene regulatory network could amplify variation from any molecular mediator in a characteristic manner once the variation is experienced within the variable genetic circuit.

The findings here may also contribute to unresolved questions in embryogenesis. The emergence of distinct cell fates from seemingly equivalent blastomeres in the mammalian embryo has been conceptualized in two ways (Chen et al., 2018). In the first, the cells of the early embryo are essentially identical blastomeres and cell fate emerges randomly (Kurotaki et al., 2007; Motosugi et al., 2005; Solter, 2016). In the second, small differences exist cell-to-cell that make fate predictable for individual blastomeres, though cells still retain a high degree of plasticity (Gardner, 2001; Goolam et al., 2016; Ju et al., 2017; Piotrowska-Nitsche et al., 2005; Torres-Padilla et al., 2007; White et al., 2016). In our view, the findings here point towards a model reconciling these views. Cells are “identical” in that each experiences inherent fluctuations at a specific subset of gene regulatory network elements. If measured precisely enough, this variation is detected because cells are not perfectly synchronized with respect to these fluctuations. It is precisely the systems level organization of variable elements into a feedback loop that allows for intrinsic heterogeneity, or plasticity between states. In other words, though states appear to emerge in a random manner from a group of nearly equivalent cells, which states emerge is deterministic, dependent on the identity of the enhancers and developmental regulators prone to variation and wired together at that point in developmental time. Therefore, the amplification of inherent fluctuations by a gene regulatory network provides a mechanism by which stereotyped states can emerge from otherwise equivalent cells in the absence of external information. The organization of fluctuations by the gene regulatory network also confers robustness on development, as it leads to particular states that can transition into each other such as observed here. The formation of new states through a feedback loop for variation could represent a recurring motif in development. Future studies will be necessary to determine if this view applies more broadly to mammalian development. Nevertheless, the results presented here suggest that naturally arising cell-to-cell variation, often described as stochastic fluctuation, is in fact coherently organized biology.

## Supporting information

Supplemental Figures and Legends (Supplemental Item 1)

Supplemental Item 2 - Primers for CRISPR/Cas9 Targeting and Genotyping

Supplemental Item 3 - Gene expression in states

Supplemental Item 4 - miRNA expression in states

## Acknowledgements

The authors would like to thank Paige Coles for technical assistance. Additionally, we thank the Koch Institute Swanson Biotechnology Center for technical support, specifically the Flow Cytometry Core and Shanu Metha, Stuart Levine, and Vincent Butty for assistance with single cell RNA-seq. SG acknowledges funding from a Charles W. and Jennifer C. Johnson Clinical Investigator Award and NIH NCI T32 CA009216, PAS acknowledges funding from NIH P01 CA042063, and CB/MdG/AdM acknowledge funding from Horizon 2020 MSC-RISE-2016 grant agreement no. 734439 (INFERNET). This work was supported in part by the Koch Institute Support (core) Grant P30-CA14051 from the National Cancer Institute.

## Author Contributions

SG conceptualized the study. MC, SH, and SG performed experiments in the laboratories of PAS and SG. MdG, AdM, and CB developed the inference model. SG wrote the manuscript with input from MC and editing by PAS.

## Methods

### EXPERIMENTAL MODEL AND SUBJECT DETAILS

#### Cell lines

Unless otherwise noted, ESC lines used in this study were derived from V6.5 mouse Embryonic Stem Cells (R. Jaenisch laboratory, Whitehead Institute MIT) and were cultured feeder free as described below. TTFHAgo2 doxycycline inducible Ago2 ESC were described in (Zamudio et al., 2014). In brief, these cells were generated from AB2.2 ESC where the endogenous copies of Ago1/2/3/4 were deleted (described in (Su et al., 2009)) and human Ago2 was re-expressed under tight doxycycline inducible control. DGCR8^−/−^ ESC are derived from V6.5 ESC (described in (Wang et al., 2007)). *Mir182*^*indel*^ and *Mir708*^*indel*^ ESC were derived from V6.5 ESC (this study) by CRISPR-Cas9 targeting of the hairpin loop for these genes (see Method Details below). All of these cell lines were also labelled at *Nanog* and *Sox2* loci (this study, see “Fluorophore tagging of pluripotency genes” in Method Details).

#### Cell line maintenance

Cells were maintained in serum + LIF culture medium on 10cm tissue culture plates pre-coated with 0.2% gelatin in phosphate-buffered saline (PBS). Plates were maintained in a humidified 5% CO_2_ incubator at 37 degrees C. Culture medium consisted of: 415 mL DMEM, 5 mL 1M HEPES, 5 mL 0.1 mM non-essential amino acids, 5 mL 0.1 mM P/S antibiotics, 5 mL 0.1 mM L-glutamine, 4 μL 14.3 M beta-mercaptoethanol, mL HyClone fetal bovine serum (FBS), and 55 μL 1000U/mL leukemia inhibitory factor (LIF). All components were passed through a sterile filter prior to addition of serum + LIF, which were added after filtration. Cells were passaged at a minimum of every two days, as follows. First, they were washed with HEPES-buffered saline (HBS); 0.5 mL of HBS was used per 1 mL of culture medium. Next, ESC were detached from the plate by trypsinization. 0.05-0.1 mL of 0.25% trypsin was used per 1 mL of medium used to maintain the cells, and trypsin was inactivated using complete culture medium after 1-2 minutes of incubation. Detached cells were spun in a centrifuge at 233 rcf for 5 minutes, and the cell pellet was resuspended in culture medium. Cells were then counted using Trypan Blue staining with a hemocytometer and re-plated at a density of 25,000-50,000 cells / mL medium or analyzed as described below.

### METHOD DETAILS

#### Theory Note – Generation of Model in Fig. 5A

Our model addresses the role of microRNAs (miRNAs) in regulating the heterogeneity of the cell states distinguished by the expression levels of Sox2 and Nanog. We consider a system formed by two targets, *R*_1_ and *R*_2_, and two distinct miRNA pools, denoted respectively by *I* and *J*. Pool *I* only contains miRNA species 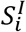(*i* indexing species in pool *I*, with *i* = 1,…, *N*) that target *R*_1_ specifically, while each miRNA species 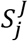 from pool *J* (indexed by *j* = 1,…, *M*) can bind both *R*_1_ and *R*_2_. In such a scheme, *R*_1_ and *R*_2_ represent respectively Nanog and Sox2.

We assume that target and miRNA abundances vary in time following:

1. transcription events,
2. degradation events, and
3. molecular titration events due to miRNA-target interactions, which after target repression, lead to miRNA degradation with probability *α*.

Each of these processes is inherently stochastic (meaning subject to inherent fluctuations in time) and occurs with a certain probability per unit time (rate). We denote by *k*_*x*_ (respectively, *g*_*x*_) the synthesis rate (respectively, the degradation rate) of species *X*, while 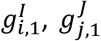 and 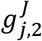represent the different miRNA-target binding rates (for instance, 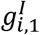stands for the binding rate between target *R*_1_ and miRNA species *i* from pool *I*). In summary, our model includes the following reactions:

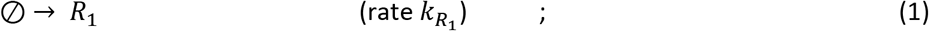

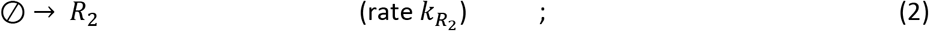

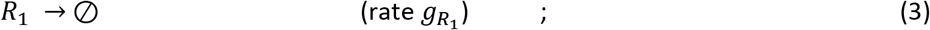

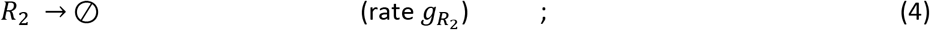

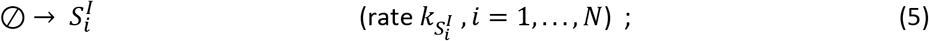

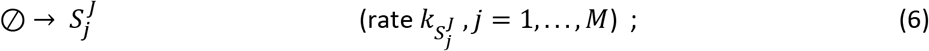

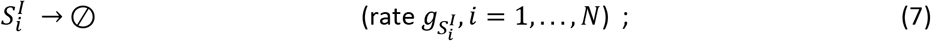

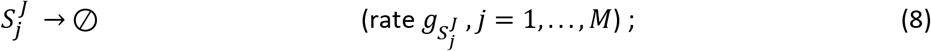

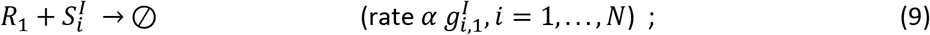

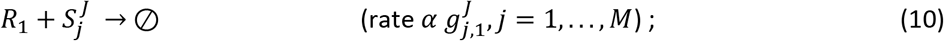

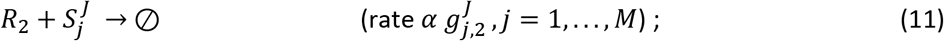

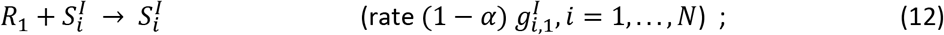

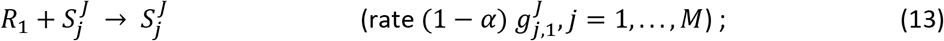

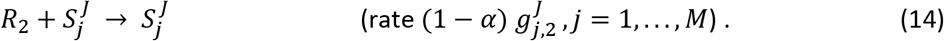

Assuming the system to be well mixed, for any fixed choice of these rates and of the parameter α, then target and miRNA abundances will change over time by fluctuating randomly around mean values described by the mass-action kinetic equations:

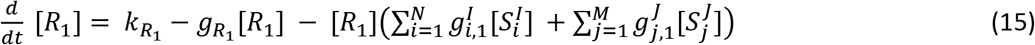

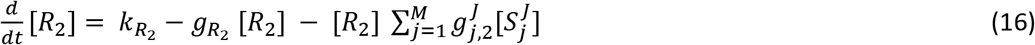

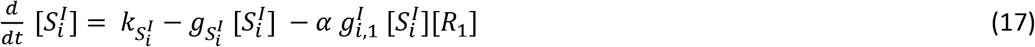

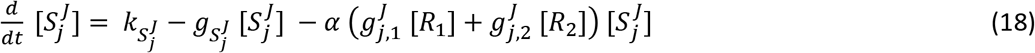

where [*X*] denotes the abundance of species *X*. At stationarity, molecular populations will fluctuate randomly around the steady state values obtained by setting time derivatives in (15)–(18) to zero and solving for [*R*_1_], [*R*_2_], 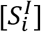 and 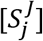. Such fluctuations can be described as intrinsic variability and arise from the inherent randomness of molecular events at given (fixed) rates.

The above setting describes how molecular levels change over time in a single cellular sample characterized by the given kinetic rates. Reaction networks like (1)–(14) can be simulated via the Gillespie algorithm (Gillespie, 1976), a broadly used Monte Carlo scheme that exploits the well-mixing assumption to conveniently schedule processes and update molecular populations over time. The Gillespie algorithm has been already applied for the analysis of miRNA-target interactions (see (Bosia et al., 2013; Martirosyan et al., 2016; Noorbakhsh et al., 2013)) and we have utilized it for studying this model *in silico*.

Besides the inherent variability due to the probabilistic nature of synthesis, degradation and interaction events, molecular levels can fluctuate cell-to-cell due to heterogeneities in kinetic rates. Such fluctuations can be termed extrinsic variability. In particular, when one considers a population of cells, the fluctuations of target levels at stationarity will generically consist of two components: one due to the intrinsic randomness of molecular events within each cells and one due to the extrinsic variability of kinetic parameters across cells. The total variance of molecular levels will therefore be given by

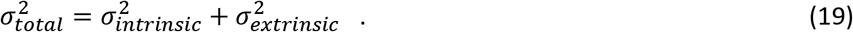

Extrinsic variability is crucial for modeling ensembles of cells. In practice, this can be achieved by simulating a large number of systems of the form (1)–(14), each characterized by different values for the kinetic rates.

To focus specifically on the contribution of miRNAs to the establishment of cell state heterogeneity, we have considered only one source of extrinsic variability, namely that affecting miRNA transcription rates. In other terms, different cells were assumed to share all kinetic rates except for miRNA transcription rates, which were taken to be different across different cells. Because of the threshold-like responses induced in targets upon increasing miRNA levels (Mukherji et al., 2011), targets can be extremely sensitive to miRNA abundances close to the equimolarity condition. This implies that when miRNA transcription rates for different cells are sampled around this threshold due to cell-to-cell heterogeneity, one will typically find the target to be expressed in some cells (at different levels) and repressed in others. Such a setup has been shown to robustly generate bimodal expression profiles even in presence of weak miRNA-RNA interactions (Del Giudice et al., 2018).

We have used the Gillespie algorithm to simulate an ensemble of miRNA-target networks described by (1)–(14), with *N* = *M* = 2 miRNA species in each pool. For each network, we fixed all kinetic parameters to representative values ensuring realistic molecular abundances and weak miRNA-target coupling (see Theory Note Table), except for miRNA transcription rates, which were randomly and independently generated for every system in the ensemble, thereby providing extrinsic fluctuations. Each network constructed in this way corresponds in practice to a cellular sample. We assumed a distribution of miRNA transcription rates of the form

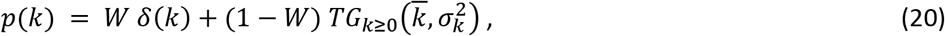

where *W* is a numerical parameter (0 ≤ *W* ≤ 1), *δ*(*k*) denotes the Dirac delta distribution, while 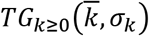 stands for the truncated Gaussian distribution defined for *k* ≥ 0, with mean 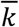 and standard deviation *σ*_*k*_. In other terms, miRNA transcription rates were taken to be nil with probability *W*, while they were sampled from the truncated Gaussian distribution with probability 1 − *W*. Ultimately, the values that *W*, 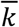 and *σ*_*k*_ take for different miRNA species define the extrinsic variability. Such values are detailed in the Table below, along with those of the remaining parameters.

We have simulated 10,000 such WT networks (each representing that of a distinct cell), probing for each system the expression levels of the targets at a time well into the stationary regime. This yields, for each sample, a point in the plane spanned by the values of *R*_1_(Nanog) and *R*_2_ (Sox2). Networks depleted of a single microRNA from the shared pool (KO or “indel” network) were simulated by simply silencing the kinetic rates of the knockout process (indicated by an asterisk in the Theory Note Table). These simulations lead to the results displayed the Theory Note Figure below.

| Parameter | Symbol [units] | Value |
| --- | --- | --- |
| Number of miRNA species in pools $I$ and $J$ | $N, M$ | 2, 2 |
| Target transcription rates | $k_{R_1}, k_{R_2}$ [molecules/sec] | 0.3, 1 |
| Target degradation rates | $g_{R_1}, g_{R_2}$ [1/sec] | 0.0009, 0.003 |
| miRNA degradation rates | $g_{S_i^I}, g_{S_j^J}$ [1/sec] | 0.0002 ($i = 1, 2$),<br>0.0002 ($j = 1, 2$) |
| miRNA-target association rates | $g_{i,1}^I$ [1/molecules·sec] | 0.000001 ($i = 1, 2$) |
| miRNA-target association rates | $g_{1,1}^J, g_{1,2}^J$ [1/molecules·sec] | 0.00001,<br>0.000003 |
| miRNA-target association rates | $g_{2,1}^J, g_{2,2}^J$ [1/molecules·sec] | 0.00005,<br>0.00008 |
| Stoichiometricity ratio | $\alpha$ (adimensional) | 0.5 |
| Prob($k = 0$)for miRNAs in pool $I$ | $W_i^I$ (adimensional) | 0.1 ($i = 1, 2$) |
| Prob($k = 0$)for miRNAs in pool $J$ | $W_j^J$ (adimensional) | 0.3 ($j = 1$),<br>0.2 ($j = 2$) |
| Truncated Gaussians: means | $\overline{k_{S_i^I}}, \overline{k_{S_j^J}}$ [molecules/sec] | -0.1 ($i = 1$)<br>-0.4 ($i = 2$)<br>-0.4 ($j = 1$)<br>0.05 ($j = 2$) |
| Truncated Gaussians: standard deviations | $\sigma_{k_{S_i^I}}, \sigma_{k_{S_j^J}}$ [molecules/sec] | 0.3 ($i = 1$)<br>0.3 ($i = 2$)<br>0.8 ($j = 1$)<br>0.11 ($j = 2$) |

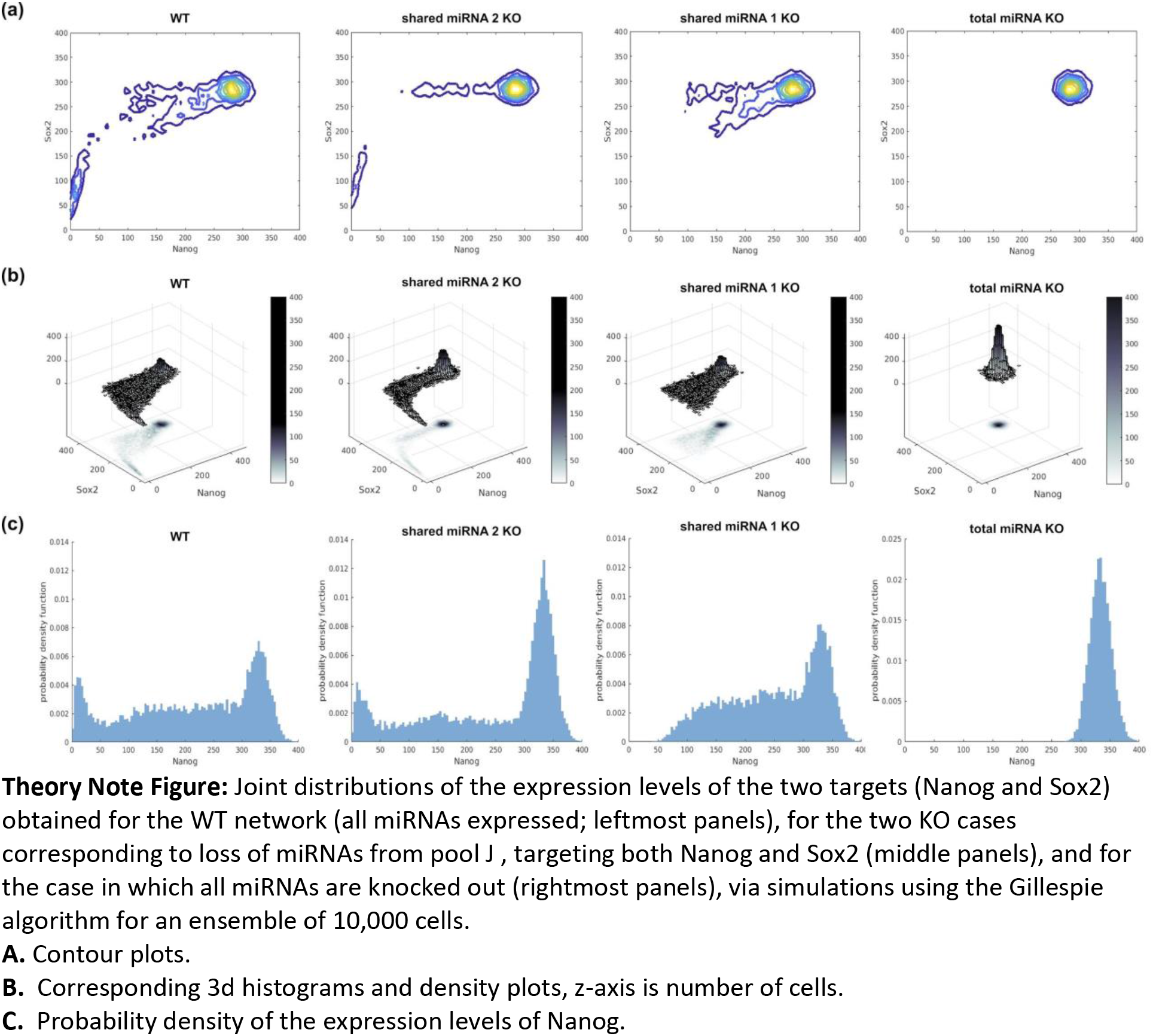

##### Fluorophore tagging of pluripotency genes

Cell lines were derived from V6.5 ESCs by inserting fluorophore tags at the endogenous loci of pluripotency genes. These cell lines are: *Nanog-GFP/Sox2-mCherry, Nanog-GFP/Esrrb-E2-Crimson, Nanog-GFP/Eif2s2-mCherry, Nanog-GFP/Hsp90ab1-mCherry*. All tagging was accomplished via CRISPR-Cas9 induced homology directed repair, using a guide RNA targeting immediately upstream of the start codon (*Nanog*) or downstream of the stop codon (*Sox2, Esrrb, Eif2s2, Hsp90ab1*) and a repair template encoding both the fluorescent marker and drug resistance separated by post-translational cleavage sequences P2A or T2A (See Fig. S1) following the method of (Stewart-Ornstein and Lahav, 2016). For each targeted gene, the guide RNA sequence (Supplemental Item 2) was cloned into the PX330 plasmid (Addgene #42230), which expresses S. pyogenes Cas9 (SpCas9) nuclease using BbsI digestion. The modified PX330 plasmid was then introduced into cells by cationic lipid transfection (Lipofectamine 2000, Invitrogen) along with a homology directed repair construct containing the relevant fluorophore, T2A/P2A, and drug resistance (see Fig. S1). Transfected cells were selected by drug resistance; PCR primers flanking the ends of the homology directing regions were used to confirm insertion of the repair construct at the endogenous locus (Supplemental Item 2).

##### Flow cytometry and fluorescence activated cell sorting (FACS)

ESCs were analyzed on either LSRII or LSRFortessa analyzers (Becton Dickinson) with FACSDiva v8.0 acquisition software. In general, cells were 50-90% confluent at time of analysis. FCS files were exported and analyzed with FlowJo V9.9 software. In brief, samples were gated first based on FSC-A vs. SSC-A scatter profiles for live cells and then based on FSC-W vs. FSC-H scatter profiles to eliminate aggregates. Cells singly transfected with transient fluorophore expression constructs or singly tagged with either GFP or mCherry were used as fluorescence compensation controls. Channel gains were adjusted based on native V6.5 ESC and adjusted as appropriate. See also “Bidirectional reporters for miRNA activity” below for additional details regarding analysis of data depicted in Fig S4C.

Sorting of *Nanog-GFP/Sox2-mCherry* ESCs into states (as defined by GFP/mCherry protein levels) was done using a FACSARIA cell sorter (Becton Dickinson). States were sorted into fresh culture medium in 5 mL collection tubes, then immediately spun at 1,000 rpm for 5 minutes and resuspended for either plating or RNA isolation (See also “ESC culture” and “RNA-sequencing sample preparation"). Single-cell sorting was done on the same FACSARIA machine. Single cells were sorted into 200 μL fresh culture medium in one well of a 96-well flat bottom plate (VWR Catalog #29442-054).

##### Molecular Barcoding of ESC

ESC were barcoded using the ClonTracer library (Addgene #67267) described previously (Bhang et al., 2015). In brief, lentivirally encoded barcodes were transduced into ESC fluorophore tagged at the *Nanog* and *Sox2* locus (*GFP-Nanog, Sox2-mCerulean3*) using spinoculation and selected by FACS sorting for RFP expression. Transduced library cells were cultured together in 10 cm plates (Fig S1) or separately as single cell clones in 96-well plates (Fig 1B) for the indicated time periods. Pooled genomic DNA was extracted using Sigma GenElute Mammalian Genomic DNA Prep Kit (Catalog #G1N70) and PCR amplified according to the protocol provided by (Bhang et al., 2015) on the Addgene website. Amplicons were sequenced by Illumina FlowCell for pools or by Sanger sequencing for single cell clones.

##### RNA-sequencing sample preparation

After separation by flow cytometric sorting, ≥ 100K cells each from States 1/2/3 or the total unseparated ESC population passed through the sorter were placed separately into 750 μL TRIzol (Thermo, 15596026) and total RNA isolated according to the manufacturer‘s protocol. Samples were treated with DNase I (NEB M0303) and RNA isolated by ethanol precipitation. Total RNA was analyzed by Agilent BioAnalyzer and accepted for sample RIN > 7.0. RNA-sequencing represents three biological replicates isolated by flow-sorting on three separate days. rRNA-depleted RNA-sequencing libraries (~100 ng RNA/sample) were prepared using Kapa RNA HyperPrep Kit with RiboErase (HMR) KK8561 using 11 rounds of PCR amplification after addition of ERCC spike-in controls at the recommended concentration. The final libraries were QC checked by fragment electrophoresis and qPCR for colony forming units prior to pooling and loading on an Illumina FlowCell (NextSeq 500, 150 bp PE reads). Each sample library was sequenced to depth of 30-45M reads.

##### RNA-sequencing analysis pipeline and data plotting

RNA-sequencing reads were first trimmed for adaptor sequence using Trimmomatic v0.36 using the following command:

~~~
java -jar $dir/trimmomatic-0.36.jar PE -phred33 $file1.fastq $file2.fastq $file1$clip.fastq
$file1$unpaired.fastq $file2$clip.fastq $file2$unpaired.fastq ILLUMINACLIP:$adap/TruSeq3-PE-2.fa:2:30:10:4:TRUE SLIDINGWINDOW:4:10 MINLEN:16
~~~

where $file1 and $file2 represent the paired-end read fastq files. Next, RNA-sequencing reads were aligned to the mm10 genome using Gencode M15 transcript annotations (www.gencodegenes.org/mouse/release_M15.html) using the comprehensive gene annotation file. Genome indices were generated for STAR (v2.4.1d) use as the aligner called by RSEM (v1.2.30) using the following command:

~~~
rsem-prepare-reference -p 8 --star --gtf $dir/gencode.vM15.primary_assembly.annotation.gtf
/home/salilg/rsem_mm10/GRCm38_M15
~~~

RSEM counts were obtained using (for example on library 3680):

~~~
rsem-calculate-expression --forward-prob 0 -p 8 --star --paired-end $dir/3680_1_sequence_clip.fastq
$dir/3680_2_sequence_clip.fastq /home/salilg/rsem_mm10/GRCm38_M15
$dir/rsem_GencodeM15/rsem_GencodeM15_3680_output_clip_genes.results
~~~

Which were then combined across all 3 replicates of each state and total population sequencing files by (where each .results file is noted by brackets):

~~~
rsem-generate-data-matrix [State 3 rep1] [State 3 rep2] [State 3 rep3] [State 2 rep1] [State 2 rep2]
[State 2 rep3] [State 1 rep 1] [State 1 rep2] [State 1 rep3] >
Sox2E12_GencodeM15_CombinedRepsinorder_clip.counts.matrix
~~~

and then inputted into EBseq‘s multi-test function using default parameters by:

~~~
rsem-run-ebseq $dir/Sox2E12_GencodeM15_CombinedRepsinorder_clip.counts.matrix 3,3,3,3
$dir/Sox2E12_GencodeM15_Combinedreps_clip_EBseq.results
~~~

Differentially expressed genes between all States were determined as those with posterior probability of differential expression (PPDE) ≥ 0.95 and fitting pattern 14 (1,2,3,3) or pattern 15 (1,2,3,4) as given by EBseq‘s multitest function for 4 distinct conditions (State3, State2, State1, All) over three biological replicates of sorting. Differentially expressed genes between any 2 States were determined as PPDE ≥ 0.95 and best matching patterns 2-13. For Figure 1C (MA plots), x-axis represents Log_10_(mean RSEM expected counts across all three States) and is displayed starting at a value of 10 (unless otherwise noted, all data analyzed in this study were filtered for RSEM expected counts > 10). For CV-mean plots, coefficient of variation (CV) was determined as: ((standard deviation of RSEM expected counts (conditional means) across all three states) / mean of RSEM expected counts averaged across all three states) and plotted against mean of RSEM expected counts averaged across all three states. Scatter plots were generated using GraphPad Prism v6.07 and cumulative density plots using ggplot2‘s geom_density() function in R. Heatmap in Fig. 1E is generated using R‘s ComplexHeatmap package (Gu et al., 2016) using RSEM‘s raw expected counts for each displayed gene in each state and replicate as follows: z-score = (Log_10_(expected count) - Log_10_(mean expected counts across all 9 state samples for this gene)) / (population standard deviation for all genes in this heatmap). For distance heatmaps, we compared expression of the top300 protein coding genes in each state that were highest expressed in that state to the expression of these same genes in mouse blastocyst (Boroviak et al., 2015; Shahbazi et al., 2017) using the manhattan distance method in R (ComplexHeatMap). We then plotted the distance matrix between conditions as a heatmap (Fig. 1G and Fig. S2D).

##### Gene Ontology Analysis

Lists of combined PPDE ≥ 0.95 pattern 14 and pattern 15 genes were inputted into GOrilla (http://cbl-gorilla.cs.technion.ac.il/) on 1-11-18 to determine ontologies highest represented in this group. Display in Fig. 1 corresponds to reduction of redundant terms. Fig. S2B displays CV-mean plot for selected gene ontologies (GO: 0048646 and 0051726) chosen to represent lineage regulators and cell cycle genes respectively of similar set size and expression distribution. To determine ontology representations of each state (Fig S2C), we took the top 300 expressed protein coding genes in each state that were highest expressed in that state and analyzed by GOrilla. Fig. S2C represents z-score of FDR q-value for selected ontology terms in each state chosen as representatives of the results. Selected terms from amongst the strongest processes for each state are shown in Table 1.

##### Enhancer Activity in ESC States

In order to ensure compatibility with previous annotations of enhancers in ESC (Dowen et al., 2014; Suzuki et al., 2017; Whyte et al., 2013), we aligned our rRNA-depleted RNA-sequencing data to the mm9 genome using STAR (v2.4.1d):

~~~
STAR --runMode alignReads --runThreadN 8 --genomeDir $dir1 -readFilesIn
$dir2/3680_1_sequence_clip.fastq $dir2/3680_2_sequence_clip.fastq --twopassMode Basic --
sjdbOverhang 149 --outFileNamePrefix $dir2/3680_STARalignment_050817 --outReadsUnmapped Fastx
--outSAMtype BAM SortedByCoordinate --outFilterMultimapNmax 20 --outFilterMismatchNmax 999
-- outFilterMismatchNoverLmax 0.1 --alignIntronMin 70 --alignIntronMax 500000 -alignMatesGapMax
500000 --alignSJoverhangMin 8 --alignSJDBoverhangMin 1 --outFilterType BySJout
~~~

Counts against enhancer features such as OSN-TE & OSN-SE or H3K27Ac-TE & H3K27Ac-were generated by taking enhancer feature annotations defined in mm9 genome coordinates in previous studies [OSN-TE and OSN-SE were given by (Whyte et al., 2013) and H3K27Ac-SE and H3K27Ac-TE were given by (Suzuki et al., 2017)] as browser extensible data (BED) format files using BEDtools v2.20.1:

~~~
bedtools multicov -D -bams [State 3 rep1 bam from above] [State 3 rep2] [State 3 rep3] [State 2 rep1]
[State 2 rep2] [State 2 rep3] [State 1 rep 1] [State 1 rep2] [State 1 rep3] [Total rep1] [Total rep2] [Total rep3] -bed [enhancer annotation] > counted_[enhancer type].bed
~~~

These counts were then prepared into a matrix file appropriate for EBseq and differential enhancer RNA production analysis using EBseq multitest:

~~~
rsem-run-ebseq $dir/counted_[enhancer type].bed 3,3,3,3 $dir/[enhancer type]_mm9_results
~~~

Similar to analysis for protein coding genes, enhancers were noted as differentially expressed between all three States for PPDE ≥ 0.95 and pattern 14 or pattern 15 assignment and between two States for pattern 2-13 assignment by EBseq. As noted above for protein coding genes, scatter plots and stacked bar graphs (Fig. 2A-B) were made using GraphPad Prism and cumulative density plots (Fig. S2B) using R.

Enhancer-gene regulation pairs were taken from ChIA-PET analysis (Dowen et al., 2014). For all enhancer annotations, a gene was determined to be regulated by that enhancer if both appeared completely within a single interval loop, the boundaries of which were defined by cohesin binding. In the case of miRNA genes (Fig. 3C), each was mapped to the nearest super enhancer (H3K27Ac) based on distance (1 Mb window) due to the paucity of SE-miRNA gene captured intervals in current ChIA-PET datasets.

##### Enhancer Reporters

Reporters of enhancer activity were constructed by cloning *mCerulean3* (Addgene #54730) into the multiple cloning site of Open Biosystems vector PB533A-2. This vector contains an internal ribosomal entry site (IRES) followed by neomycin resistance followed by SV40 polyA, and this entire fragment including the preceding EF-1α promoter was amplified by PCR (Forward 5‘-GGGCAGAGCGCACATCG-3’ and Reverse Primer 5‘-CAGACATGATAAGATACATTGATGAGTTTGG-3’) The linear PCR product was then gel purified and inserted into enhancers for *Nanog*, *Pou5f1*, *Esrrb*, and *Fgf4* using non-homologous end joining (NHEJ repair) induced by CRISPR-Cas9 targeting adjacent to the enhancer (see Supplemental Item 2 for Cas9 guide sequences). Cells were selected in G418 300 μg/mL for 5 days, single cell cloned by sorting, and grown for ~10 days. These clones were then genotyped for proper insertion of the reporter by PCR using primers 50-100 bp upstream and downstream from the CRISPR-Cas9 cutsite (Supplemental Item 2) and sequencing PCR bands. We selected clones containing >50 bp residual genomic sequence along with full insert sequences (EF-1α promoter-*Cerulean*-IRES-*neo*-polyA) at the band consistent with insertion size. Additionally, we chose clones with relatively unperturbed distributions of Nanog vs Sox2 to indicate cell state. At several additional enhancer sites aside from those indicated by the sgRNA sequences given (Supplemental Item 2), no clones were identified with intact cell state distributions or with insertions containing intact sequences and these were not included in our study (data not shown). Reporter activity along with cell state distributions was then measured by flow cytometry for Cerulean, GFP (Nanog), and mCherry (Sox2) using singly labelled cells as compensation controls on a Beckton-Dickinson LSR Fortessa instrument using 405, 488, and 561 laser lines. For analysis, using FlowJo v9.9, cells were gated on live singlets by FSC-A/SSC-A/FSC-W/FSC-H, and were either further gated on States (GFP vs mCherry, Fig. 2C) or the data for all 3 fluorophores exported as tables and plotted using plot.ly for Python (Fig. 2D).

##### Metagene Plots

Metagene binding plots (Fig. 2G) were generated by analyzing ChIP-seq binding profiles of Nanog, Sox2, Esrrb, Pou5f1, Smad1, and Tcf3 in embryonic stem cells and comparing them to input. Files utilized were as follows: SRR713342 (Nanog), SRR713341 (Sox2), SRR001992 (Esrrb), SRR713340 (Pou5f1) normalized to input SRR713343 and SRR002020 (Smad1), SRR015155 (Tcf3) normalized to input SRR058997. These data derive from (Hnisz et al., 2013; Whyte et al., 2013) and were downloaded from the Gene Expression Omnibus and mapped to the mm9 genome using bowtie as described (Suzuki et al., 2017). “.bam” files were loaded into the R package metagene (Beauparlant CJ, 2019) and tested against OSN-SE regions given by (Dowen et al., 2014) using the command metagene$new(regions = regionOSN, bam_files = bam_file_vector, padding_size = 500) which extends each enhancer region 500 bp on either side. Binding in these regions was then given in 1000 bins by using $produce_table(design=design, normalization = ‘RPM‘, bin_count=1000) and the middle 400 bins plotted as shown.

##### Single cell RNA-sequencing and gene neighborhood construction

Single cell RNA-sequencing (scRNA-seq) was performed by the Koch Institute Nanowell Cytometry Core using SeqWell (Gierahn et al., 2017) technology. In brief, single cell suspensions of fluorophore tagged V6.5 ESC (cultured in serum + LIF) were made by trypsinization followed by serial passage through 50 micron cell strainer meshes. Approximately 10,000 cells were loaded onto a SeqWell array, lysed, and prepared as single cell cDNA libraries as described (Gierahn et al., 2017). Libraries were sequenced using a NextSeq 500 and aligned to the mm10 genome using the SeqWell analysis pipeline generated by the C. Love Lab at MIT. The entire process was repeated using a second preparation of ESC on a different day (and different array/sequencing run) to generate two data tables representing reads per cell across mm10 annotated transcripts (Gencode M15). These two tables were merged together and analyzed. To ensure robust inferred neighborhoods based on correlation, cells with fewer than 5,000 uniquely captured transcripts were discarded from analysis yielding 2,299 cells remaining. We then restricted our analysis to the best sampled genes, keeping those with a total of >500 counts across all remaining cells OR with >4 counts in any one cell to account for rare cells with high expression of an individual transcript. Additionally, analysis comparing WT ESC and *Mir182*^*indel*^ ESC are restricted to 6,107 genes captured by these criteria in both cell types. Data tables were then processed by total count normalization (Klein et al., 2015) prior to neighborhood analysis. Raw fastq files, count tables and merged count tables, and normalized tables are provided.

To construct interaction neighborhoods, we utilized a topology based method similar to (Klein et al., 2015) reasoning similarly to these authors that such a method was most likely to minimize artifacts from technical sampling noise in scRNA-seq. This method is based originally on the work of (Li and Horvath, 2007). In brief, a node gene of interest is selected (G_0_ gene). The 50 most correlated (Pearson r correlation coefficient) genes with this node are then selected (G_1_ set), and then the 50 most correlated with each of these G1 genes is selected (50 G_2_ sets for every G_1_ set). Thus an initial network is created with 2550 directed edges (G_0_->G_1_ or G_1_->G_2_), many of which represent mutual relationships between genes (e.g. two distinct G_1_ genes can be in each other’s G_2_ sets). We then iteratively trim the network by removing any genes with < 10 incoming edges (from G_0_ or G_1_ genes) including removing any outgoing edges for removed G_1_ genes until we are left with a final network that must include G_0_ and in which all genes have ≥ 10 incoming edges. We found relatively little difference in final networks when choosing 30-75 for the initial number of correlated genes in each G_1_/G_2_ set or network trimming requiring 5-12 incoming edges.

##### Single Cell Variation (ν) Score Calculation

We calculated a “variability” score to describe the variation in the expression of a gene across a population of cells, based on a test statistic described in (Klein et al., 2015). For any given mean expression level, this test statistic weights genes whose coefficient of variation is significantly larger than a Poisson random variable with the same mean. The test statistic, ν, is:

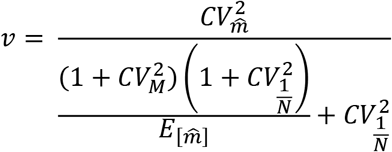

where 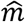 is normalized read counts (total count normalization), M is total number of reads and N is total mRNA content. The additive constant noise 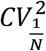 weights against genes whose variation in expression is largely due to variation in differences in cell size. In this study, we use 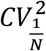 of 0.25.

##### Single Cell State Analysis (PHATE mapping, PCA, Force Directed Graphs)

Single Cell state representations were made primarily using the PHATE (Moon et al., 2019) and ScanPy (Wolf et al., 2018) packages installed in Python 3.7. A full script with detailed commands used to produce plots in Fig. 6 and S7 is given at: https://bit.ly/2qfUSLQin/scripts/Seurat_Python2.py. In brief, the combined (WT and *Mir182^indel^*), mapped single cell expression tables were normalized and then analyzed in PHATE using default parameters to power the PHATE operator. Cells were pruned by removing those with the top 5% of mitochondrial reads, transformed by square root, and the total number of counts and percent of mitochondrial reads regressed out using ScanPy prior to applying the PHATE operator. PCA and force directed graphs were generated using built-in ScanPy functions using default parameters in analogy to the methods shown in (Paul et al., 2015). Cells were colored according to their Z-score for State 1-3 genes in the following way. First, the top300 genes uniquely most highly expressed in each state (see GO analysis or Heatmap analysis) are defined. Then, for each gene, a Z-score for its expression across cells is calculated for each cell. Next, the Z-scores for all 300 State 1 genes are summed together for each cell, as are the Z-scores for all 300 State 2 genes and State 3 genes. These summed Z-scores are then re-scaled across all cells to the range 0,1 for each state. Thus, the cell with the ‘most State 1 character’ is given a State1Z-score of 1.0 and that with the least such character is given a score of 0. These scores were then used for red/blue coloration in Figures 6 and S7.

##### MiRNA-sequencing and data analysis pipeline

Total RNA samples were prepared from ESC states identically to RNA-sequencing above for two biological replicates isolated by flow cytometric sorting. Small RNA libraries were then prepared using the NEB small RNA-sequencing kit (E7300S) according to the manufacturer‘s instructions using 13 cycles of PCR amplification. QC assessment was done by electrophoresis and colony forming units prior to loading pooled samples onto an Illumina FlowCell (HiSeq2000, 1 lane for 8 samples, 40 bp SE reads) with 10-15M reads/sample. In order to map miRNA reads in the background of the transcriptome and avoid issues with forced mapping to an artificial genome, we inserted mature miRNA annotations (miRbase v21, www.mirBase.org) into the Gencode M15 main annotation gtf file. Mature miRNA seeds were inserted as distinct transcripts (with supporting exon annotations) with appropriate parent miRNA genes, allowing alignment and single step counting of gene and mature miRNA levels. This combined gtf annotation file is available at: https://bit.ly/2qfUSLQ.

To map small RNA-sequencing, we first generated STAR mapping of small RNA-seq to the transcriptome in the background of the mm10 genome using parameters derived from the ENCODE small RNA-seq pipeline (for sample 4284 shown as an example):

~~~
STAR --runMode alignReads --runThreadN 8 --genomeDir $genomedir -readFilesIn
$dir/4284_sequence_clip.fastq --outFileNamePrefix $dir/4284_5_STAR_mm10_042918 --
outReadsUnmapped Fastx --outSAMtype BAM SortedByCoordinate --outFilterMismatchNoverLmax 0.05
--outFilterMatchNmin 16 --outFilterScoreMinOverLread 0 --outFilterMatchNminOverLread 0 --
alignIntronMax 1 --sjdbGTFfile $genomedir/gencode.vM15.miRBase.gtf --sjdbOverhang 74 --quantMode TranscriptomeSAM
~~~

Next, we prepared RSEM reference indexes using our custom gtf file and generated counts using (shown for file 4284 for example):

~~~
rsem-calculate-expression --bam --forward-prob 0.5 -p 4 --no-bam-output -calc-pme -seed-length 16
$dir/4284_5_STAR_mm10_042918Aligned.toTranscriptome.out.bam
$genomedir/GRCm38_GencodeM15_miRBase $outputdir/4284_5
~~~

In the case of gene level miRNA plots and analysis (Fig. 3A-B), we used EBseq gene level analysis to determine differentially expressed miRNA (DE-miRNA) using rsem-run-ebseq similar to protein coding genes and enhancer activity above and considering DE-miRNA between all 3 states to be miRNA genes with PPDE ≥ 0.95 and consistent with pattern 14 or pattern 15. In the case of mature miRNAs (processed 5p or 3p arms), we utilized our gtf to perform isoform level mapping which specified counts for 5p/3p each using:

~~~
rsem-run-ebseq --ngvector $genomedir/GRCm38_GencodeM15_miRBase.ngvec
$[dir]Sox2E12_sRNAseq_GencodeM15miRBase_isoforms_CombinedRepsinorder.counts.matrix 2,2,2,2
$[dir]Sox2E12_sRNAseq_GencodeM15miRBase_isoforms_EBseq.results
~~~

Mature miRNAs with the same seed sequence were combined into seed families similar to previous analyses (Bosson et al., 2014), and the pooled seed family was called differentially expressed between states if ≥ 30% of the total seed pool originated from a differentially expressed mature miRNA ‘isoform.‘ These pooled seed miRNA families were used for analyses shown in Figs. 3D-E, 4, S4, S5.

##### miRNA target analysis

Genomic targets of miRNAs were determined primarily based on crosslinking and immunoprecipitation (iCLIP) experiments previously described (Bosson et al., 2014). In brief, this analysis identified 6,816 high confidence “clusters” of Argonaute 2 binding through comparison of miRNA bound to TTFHAgo2 cells to ‘miRNA’ bound to TTAgo2 cells immunoprecipitated with anti-HA antibody. The list of these clusters with genomic locations and nucleotide sequences is given at https://bit.ly/2qfUSLQ (“miRNA targets” → “Ago2clusters.txt”). Note that in the iCLIP technique, stacked 5P sequencing ends indicate the site of crosslinking and putative exact location of miRNA binding within each cluster (Bosson et al., 2014). In “Ago2clusters.txt,” the notation in the “Stacked 5P end” column is as follows: -_1 indicates no observed stacked 5P end; otherwise, position of the stacked 5P end, with respect to the cluster “start” location, is given, followed by an underscore and the number of reads at that position. Multiple putative stacked 5P ends are separated by semicolons prior to the underscore.

We assumed that each distinct Ago2 cluster resulted from the binding of one particular miRNA. To determine which miRNA gave rise to each cluster, we first made two assumptions: 1) if the cluster had stacked 5P end reads, it most likely arose from the activity of a miRNA with the highest affinity target site (seed match) immediately upstream of the stacked 5P end location (Bosson et al., 2014) and 2) if the cluster did not have a stacked 5P end, it arose from the activity of the highest expressed miRNA with a predicted seed match to a target site anywhere in the cluster. In brief, higher affinity ‘8-mer’ matches between mRNA target and miRNA were prioritized over lower affinity ‘6/7-mer’ matches, and higher expression miRNAs were prioritized over lower expression miRNAs when no evidence for strong affinity-based interaction was present. miRNAs with identical ‘7-mer’ seed region sequence were grouped for all analyses. Commented code for generating miRNA-mRNA assignments is available at https://bit.ly/2qfUSLQ (“Code for making biochemical miRNA-target map.py”) as is the resulting miRNA-target map (“Biochemical miRNA-target map.txt”). Note that the map only predicts targets for the first 250 miRNAs in “Information about miRNA families.txt.” These are the top 250 expressed miRNAs in the *Nanog-GFP/Sox2-mCherry* ESCs used in this study. For the analysis in Fig. S4A, we used “TargetScan miRNA-target map.txt.” This map was generated by using three tables of data (“Predicted Conserved Targets,” “Conserved Family Info,” and “Nonconserved Family Info”) from the TargetScanMouse database (v7.1 – http://www.targetscan.org/cgi-bin/targetscan/data_download.cgi?db=mmu_71), and also only predicts targets for the top 250 expressed miRNAs in the *Nanog-GFP/Sox2-mCherry* ESCs.

##### MiRNA enrichment in neighborhoods

Once we had determined an appropriate map of miRNA-mRNA interactions we utilized this for the analysis in Figs. 4, S5. To determine miRNA enrichment within a given gene network neighborhood (Network N_0_), we first listed the total number of binding events for each of the top 250 miRNA seed families within all neighbors. Next, we constructed 10,000 control neighborhoods (N_1_ set), each of which had an equal number of genes to N_0_, with each N_1_ control neighborhood also chosen to contain: roughly the same average gene expression of the same expression distribution as N_0_ (choosing the same number of genes from the lowest and highest expression quartiles as N_0_, ensuring the final control neighborhood has mean expression within 0.8 * neighborhood mean expression of N_0_ * 1.2), the same number of highly Ago2 bound genes as N_0_ (genes with >5 clusters of bound Ago2-miRNA), and roughly the same number of total miRNA binding events as N_0_ (within 0.9 * number of Ago2 clusters * 1.2). These simulated, control neighborhoods (N_1_) thus represented a closely matched group to the original gene network neighborhood under interrogation. For each of the top 250 miRNA seed families, we then empirically determined the total number of binding events in each N_1_ control neighborhood. A miRNA seed family was enriched in a N_0_ gene network neighborhood if 500 out of 10,000 N_1_ control neighborhoods or fewer (empirical p-value 0.05) contained a total number of miRNA binding sites ≥ the total number observed in the N_0_ gene network neighborhood. Given the relative rarity of genes bound by > 5 miRNAs and the stringency of these parameters, many N_1_ control neighborhoods shared considerable overlap with the N_0_ gene network neighborhood under interrogation, giving us a conservative list of enriched miRNAs for each N_0_. Next, we defined groups of genes such as pluripotency (taken from reference (Klein et al., 2015), 57 total expressed in our scRNA-seq data with non-zero Ago2 binding in their network), cytoskeleton (GO: 0007010, 192 expressed), or highly variable genes (defined by ν test-statistic > 3.0 in our scRNA-seq data, 109 genes). For each of the top 250 miRNA seed families, we totaled the number of times it was enriched in these sets of N_0_ neighborhoods and plotted (Figs. 4B, S5E) or analyzed for overlaps in enrichment between sets of N_0_ neighborhoods (Fig. S5F).

##### Bidirectional reporters for miRNA activity

Activity of individual miRNAs in ESCs was determined using a bidirectional reporter system similar to previously reported (Mukherji et al., 2011) (Fig. S4C). In brief, reporters were derived from the Takara pTRE-Tight-BI plasmid in which two fluorophores are expressed under tight co-transcriptional control under regulation by TET-on promoter system. The reporters used in this study are derived from (Bosson et al., 2014) and contain mCherry fluorescence as a readout of miRNA activity, but are modified to contain *mCerulean3 (CFP)* substituted for *YFP* due to an observed "tighter" correlation between mCerulean3 and mCherry than between YFP or ZsGreen and mCherry (see Fig. S4C). Additionally, the *mCherry* UTR contains a small insertion to introduce a SpeI restriction site for cloning. MiRNA activity is detected via insertion of target sites for the relevant miRNA into the 3’ UTR of *mCherry* using mCerulean3 as a normalization control. We chose to insert three perfect matches to each miRNA, separated by spacer sequences (cloned using ClaI and SpeI). Spacer sequences: Upstream of each miRNA site – CTGGGCACCAACTCAACTTC, Downstream of each miRNA site – ACAACTTGGTGTGTTAGTGT.

Activity of a given miRNA is determined by comparing mCherry expression in cells transiently transfected (Lipofectamine, see above) with a control plasmid (i.e., one with no miRNA sites in the *mCherry* 3’ UTR) to mCherry expression in cells transiently transfected with a plasmid with miRNA 3X sites in the 3’ UTR (additionally, cells are cotransfected with rtTA plasmid to engage the Tet system). This comparison generates the “percent silencing” for a miRNA compared to a no-sites control as follows: Percent silencing in a window of CFP expression = 100% * (1 - (mean mCherry fluorescence in cells transfected with control plasmid) / (mean mCherry fluorescence in cells transfected with plasmid with miRNA 3X sites)).

##### Generation of Mir^indel^ ESCs

MiRNA-deficient cell lines (termed *Mir*^*indel*^) were derived from the *Nanog-GFP/Sox2-mCherry* ESCs generated by fluorophore tagging of V6.5 ESCs (see “Fluorophore tagging of pluripotency genes,” above). Specifically, to achieve deficiency for a particular miRNA, CRISPR-Cas9 expressing a guide sequence targeted near the hairpin stem-loop was used to introduce an indel in the relevant miRNA gene. PX330 plasmid containing the relevant guides was transiently transfected into cells and single cell transfectants were isolated by Fluorescence-activated cell sorting (FACS) into individual wells of a 96-well flat-bottom plate. Guide RNA sequences with overhangs to facilitate BbsI cloning into pX330 are listed in Supplemental Item 2; forward and reverse sequences were annealed prior to cloning. PCR primers flanking the endogenous miRNA gene locus (Supplemental Item 2) were used to confirm the presence of an indel. Reduction in mature miRNA production was confirmed using RNA extraction by TriZOL^®^ and RT-qPCR using the Mir-X miRNA First-Strand Synthesis Kit (Catalog number 638315). Mir-X primers used for this purpose:

miR-182: TTTGGCAATGGTAGAACTCACACCG
miR-708: CAACTAGACTGTGAGCTTCTAG

##### Reintroduction of miRNAs into deficient ESCs

MiRNA re-expression in *Mir*^*indel*^ cells was performed as follows. First, bidirectional reporter constructs were generated in which *mCherry* was replaced with the *pri-miRNA* sequence. Primers used to amplify *pri-miRNA* for cloning into pTRE-BI-Tight were AATCGGATCCTCACTGCCTAATGCCCCTAC and GCAAAAGCTTAGCCATCTGTCTCTCCCTCA for miR-182 using HindIII and KpnI restriction sites (to replace *mCherry*) and AATCGGATCCTGAATAGCCAATGAAAATGACTTG and ATTGAAGCTTCAAGCCCAGGAGTTGAAGAG for miR-708 using HindIII and BamHI restriction sites (also replacing *mCherry*) for cloning into pTRE-Tight-BI. The use of the bi-directional expression plasmid allowed detection of pri-miRNA expression by CFP fluorophore expression. These vectors were then separately transfected into *Mir*^*indel*^ cells along with rtTA plasmid and the distribution of cells into States was determined by flow cytometry as described above. RNA extraction by TriZOL^®^ and RT-qPCR using the Mir-X miRNA First-Strand Synthesis Kit (Catalog number 638315) was used to confirm restoration of the relevant miRNA to at or above WT levels. The relevant miRNA sequence, with U replaced by T, was used as the forward primer for Mir-X (sequences above, same as primer used for indel confirmation).

### QUANTIFICATION AND STATISTICAL ANALYSIS

For all analyses with p values, significance was determined at p <= 0.05. P values are shown on the figure wherever they are used.

The one-sided Kolmogorov-Smirnov (K-S) test statistic was used to assess whether paired cumulative distribution functions (CDFs) were significantly different (Figs. 3D, S4A, S4D, and S4E). The one-sided test was used because in all cases, a sample distribution was being compared to a reference distribution. The Kruskal-Wallis test statistic was used to assess differences between 3+ CDFs (Figs. 2F and 3E). Both the and Kruskal-Wallis test statistics were calculated in GraphPad Prism.

The cumulative hypergeometric statistical test for enrichment was used in Figs. 3C, S3D, and S4A (right). This test detects enrichment for a property in a population sample, compared to what would have been expected based on the prevalence of that property in the whole population. Four numbers are required: population size (N), number of population successes (n), sample size (K), and number of sample successes (k). We calculated the test statistic in Python, taking advantage of scipy.stats.hypergeom.pmf and summing from k to min(n,K) to calculate the cumulative value.

Values for N, n, K and k are as follows:

3C – 315, 62, 32, 11
S3E – 724, 345, 168, 103
S4A (right) – 6721, 4403, 926, 653

Box-whisker plots were generated using plot.ly for Python (Figs. 2D and 5E) or matplotlib (Figs. 5B and 5C). All box-whisker plots use the middle 50% of the data, so the whiskers extend from the 25^th^ to the 75^th^ percentile, the middle line is drawn at the 50^th^ percentile, and the lines of the boxes are drawn at the 37.5^th^ and 62.5^th^ percentiles.

## DATA AND SOFTWARE AVAILABILITY

RNA-sequencing and small-RNA sequencing data are in the process of submission to the Gene Expression Omnibus. All data and scripts used in this study are publicly available at https://bit.ly/2qfUSLQ. We are committed to providing ready access to all materials and data used in this study.

## Supplemental Items

1. (PDF) Supplemental Figures and Legends
2. (Excel Table) Primers for CRISPR-Cas9 targeting and genotyping; related to Figures 1, 2, 5, S3, and S5
3. (Excel Table) Gene expression in states; related to Figure 1
4. (Excel Table) miRNA expression in states; related to Figure 3

