## Supplemental Figures and Legends (Supplemental Item 1) for "Networks of enhancers and microRNAs drive variation in cell states"

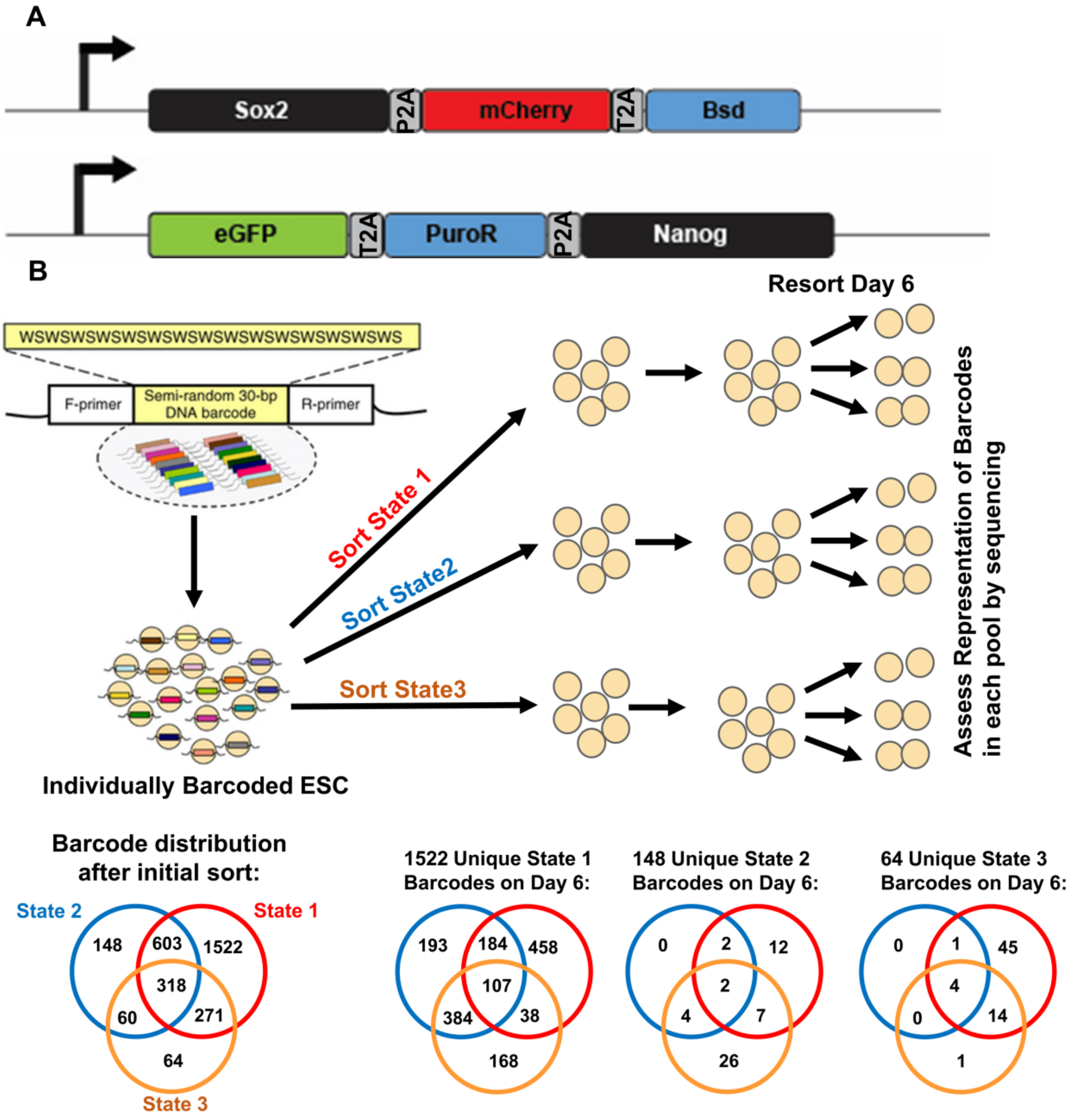
A
Sox2
P2A
mCherry
T2A
Bsd
eGFP
T2A
PuroR
P2A
Nanog
B
WSWSWSWSWSWSWSWSWSWSWSWSWSWSWS
F-primer
Semi-random 30-bp DNA barcode
R-primer
Individually Barcoded ESC
Sort State 1
Sort State2
Sort State3
Resort Day 6
Assess Representation of Barcodes in each pool by sequencing
Barcode distribution after initial sort:
State 2
State 1
State 3
148
603
1522
318
60
271
64
1522 Unique State 1 Barcodes on Day 6:
193
184
458
107
384
38
168
148 Unique State 2 Barcodes on Day 6:
0
2
12
2
4
7
26
64 Unique State 3 Barcodes on Day 6:
0
1
45
4
0
14
1

**Supplemental Figure 1: Embryonic stem cells show intrinsic heterogeneity in cell states; related to Figure 1**

**A**. Schematic depicting the modifications of *Nanog* and *Sox2* loci in embryonic stem cells used in this study. See Methods for further details on the generation of cell lines using these reporters.

**B**. A unique barcode was introduced into each ESC following the method of (Bhang et al., 2015). States 1-3 were bulk sorted, cultured for 6 days, and resorted into States 1-3. The state distribution of unique barcodes was assessed directly after initial sorting, and again after resorting on Day 6. The leftmost Venn diagram shows the number of unique barcodes in each state after initial sorting. At right are the state distributions of barcodes that initially appeared only in one state (1, 2, or 3) and were captured again on Day 6 in an unsorted pool of barcoded cells. In this way, analysis was restricted to barcode sequences present throughout the duration of the experiment (~9 days). Note that barcodes initially present only in State 2 or State 3 appear in other states on Day 6 whereas a fraction of State 1 only barcodes (458/1522) remain exclusively in this state.

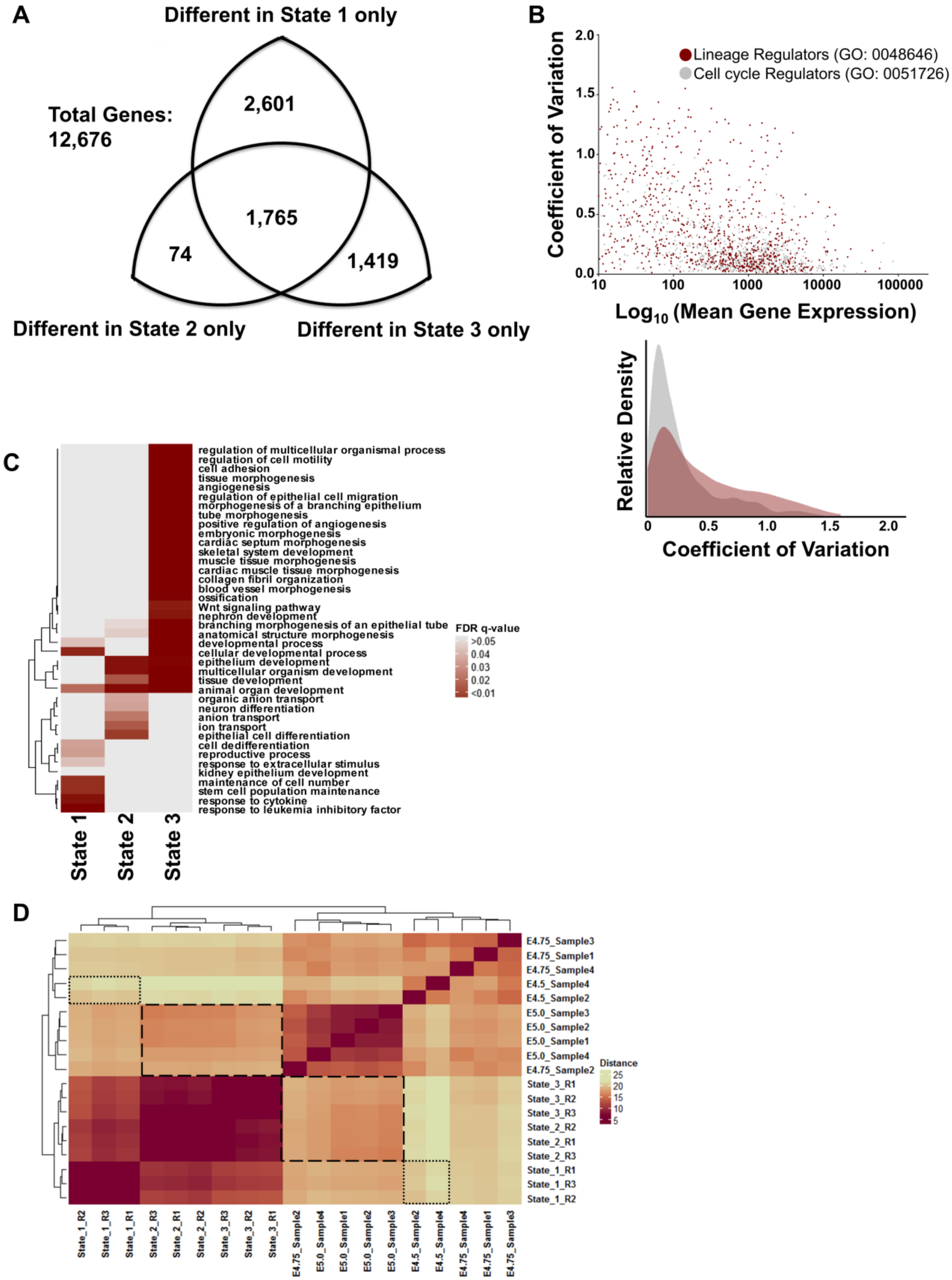
A
Different in State 1 only
Total Genes:
12,676
2,601
1,765
74
1,419
Different in State 2 only
Different in State 3 only
B
Lineage Regulators (GO: 0048646)
Cell cycle Regulators (GO: 0051726)
Coefficient of Variation
2.0
1.5
1.0
0.5
0.0
10
100
1000
10000
100000
Log10 (Mean Gene Expression)
Relative Density
0
0.5
1.0
1.5
2.0
Coefficient of Variation
C
regulation of multicellular organismal process
regulation of cell motility
cell adhesion
tissue morphogenesis
angiogenesis
regulation of epithelial cell migration
morphogenesis of a branching epithelium
tube morphogenesis
positive regulation of angiogenesis
embryonic morphogenesis
cardiac septum morphogenesis
skeletal system development
muscle tissue morphogenesis
cardiac muscle tissue morphogenesis
collagen fibril organization
blood vessel morphogenesis
ossification
Wnt signaling pathway
nephron development
branching morphogenesis of an epithelial tube
anatomical structure morphogenesis
developmental process
cellular developmental process
epithelium development
multicellular organism development
tissue development
animal organ development
organic anion transport
neuron differentiation
anion transport
ion transport
epithelial cell differentiation
cell dedifferentiation
reproductive process
response to extracellular stimulus
kidney epithelium development
maintenance of cell number
stem cell population maintenance
response to cytokine
response to leukemia inhibitory factor
FDR q-value
>0.05
0.04
0.03
0.02
<0.01
State 1
State 2
State 3
D
E4.75_Sample3
E4.75_Sample1
E4.75_Sample4
E4.5_Sample4
E4.5_Sample2
E5.0_Sample3
E5.0_Sample2
E5.0_Sample1
E5.0_Sample4
E4.75_Sample2
State_3_R1
State_3_R2
State_3_R3
State_2_R2
State_2_R1
State_2_R3
State_1_R1
State_1_R3
State_1_R2
Distance
25
20
15
10
5
State_1_R2
State_1_R3
State_1_R1
State_2_R3
State_2_R1
State_2_R2
State_3_R3
State_3_R2
State_3_R1
E4.75_Sample2
E5.0_Sample4
E5.0_Sample1
E5.0_Sample2
E5.0_Sample3
E4.5_Sample2
E4.5_Sample4
E4.75_Sample4
E4.75_Sample1
E4.75_Sample3

**Supplemental Figure 2: ESC variation between states resembles developmental programs; related to Figure 1**

**A**. Diagram of the breakdown in unique expression patterns for the 5,859 genes with significant expression differences between states (Baysian posterior probability of differential expression, PPDE ≥ 0.95). 1,765/5,859 genes showed differential expression between all three states; the remaining 4,094 genes were different in one state and the same in the other two states.

**B**. Coefficient of variation (CV) across states (y-axis) versus mean gene expression across all three states (x-axis) for lineage regulators and cell cycle genes. All genes expressed in ESC (≥ 10 expected counts) from the listed GO terms were included. Note the increased CV for lineage regulators relative to cell cycle genes. The CV data are also shown as an overlaid histogram below to facilitate comparison.

**C**. The top 300 genes uniquely highest expressed in each state were analyzed for ontology enrichment against the background of all expressed protein coding genes in ESC. Shown is a heatmap of FDR-q values for selected representative ontology terms.

**D**. Heatmap of gene expression distances between States 1-3 when compared to blastocyst expression profiles derived from days E4.5, E4.75, E5.0, and E5.5 from reference (Shahbazi et al., 2017). Note that short dashes highlight the comparison of State 1 with E4.5 samples and long dashes highlight the comparison of E5.0 with States 2-3. Comparison with all given sequencing replicates from the indicated reference are shown. See Methods for further details.

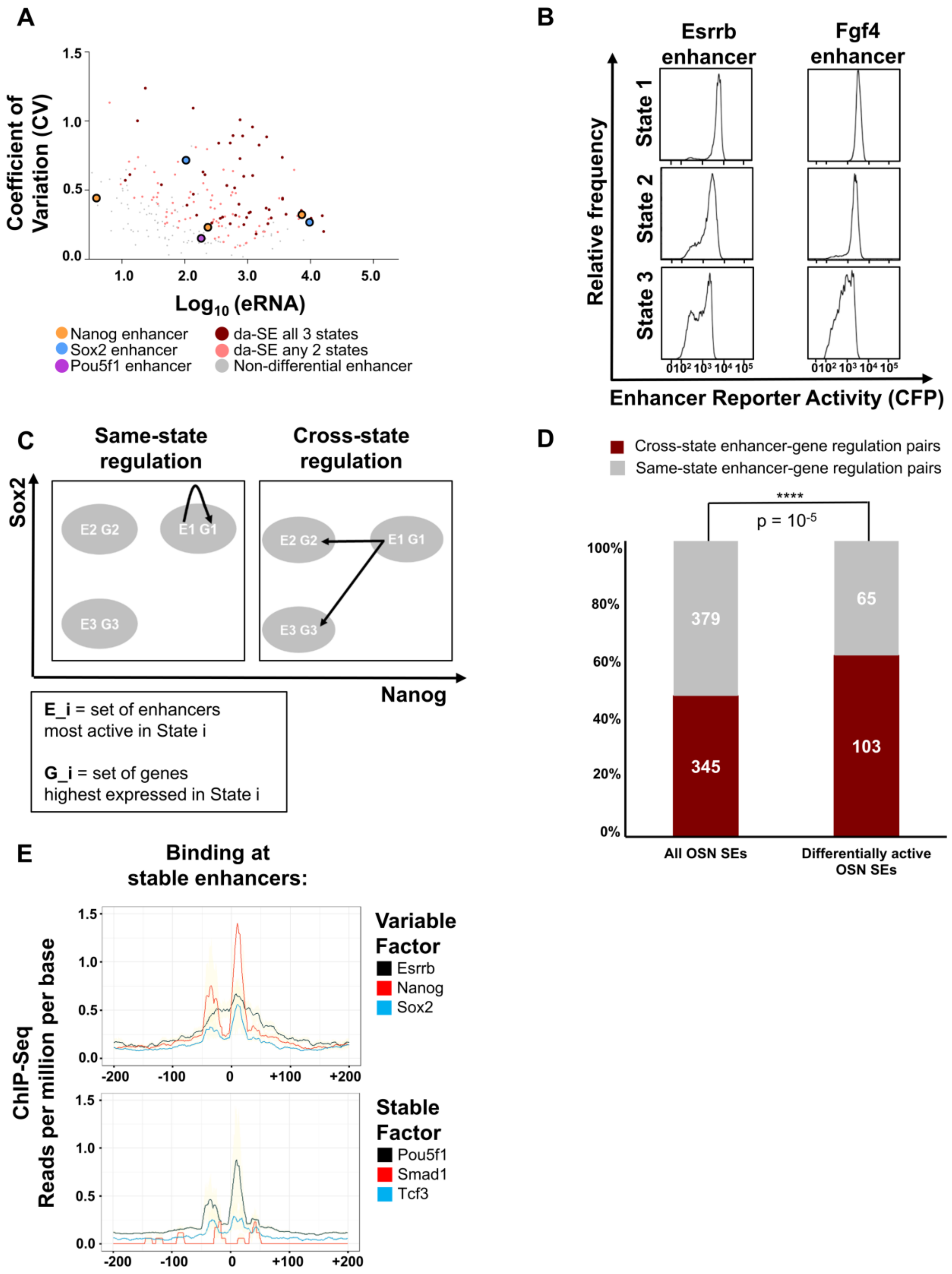

A
Coefficient of Variation (CV)
Log10 (eRNA)
Nanog enhancer
Sox2 enhancer
Pou5f1 enhancer
da-SE all 3 states
da-SE any 2 states
Non-differential enhancer
B
Esrrb enhancer
Fgf4 enhancer
State 1
State 2
State 3
Relative frequency
Enhancer Reporter Activity (CFP)
C
Same-state regulation
Cross-state regulation
Sox2
Nanog
E2 G2
E1 G1
E3 G3
E_i = set of enhancers most active in State i
G_i = set of genes highest expressed in State i
D
Cross-state enhancer-gene regulation pairs
Same-state enhancer-gene regulation pairs
****
p = 10-5
379
345
65
103
All OSN SEs
Differentially active OSN SEs
E
Binding at stable enhancers:
ChIP-Seq Reads per million per base
Variable Factor
Esrrb
Nanog
Sox2
Stable Factor
Pou5f1
Smad1
Tcf3

**Supplemental Figure 3: Properties of variable enhancers; related to Figure 2**

**A.** CV-mean plot for SE activity across three states for OSN-SEs. Highlighted in red are enhancers with significantly differential activity between all three states (PPDE ≥ 0.95, labelled da-SE). In peach are enhancers with PPDE ≥ 0.95 between any two states; in gray are other enhancers. Enhancers associated with *Nanog*, *Sox2* and *Pou5f1* are labelled.

**B**. Reporters (promoter-*CFP*-IRES-*neo*-polyA) were inserted at enhancers controlling *Esrrb* and *Fgf4* in separate cell lines (which were also labelled at *Nanog* and *Sox2* loci, see Fig. S1A). Distributions of relative reporter activity (CFP) in States 1-3 is shown.

**C**. Representation of same-state enhancer-gene regulation vs. cross-state enhancer-gene regulation. (E_i, G_j) represent enhancer-gene regulation pairs. E_i indicates the set of enhancers highest expressed in State i; G_j indicates the set of genes highest expressed in State j. Same state regulation occurs for all enhancer-gene interactions given in (Dowen et al., 2014) where i=j, and cross state regulation occurs for all pairs where i≠j.

**D**. Proportion of same-state vs. cross-state enhancer-gene regulation pairs for all OSN-SEs and differentially active OSN-SEs (colored red in S3A). A hypergeometric p-value for enrichment is shown.

**E**. ChIP-seq binding (reads per million mapped per base) at other-SEs. Each enhancer was extended to a minimum of 1 kb, all enhancers were scaled to 1000 bins, and reads normalized to input. Shown are the middle 400 bins of binding for the indicated genes. Note axes are scaled identically to allow comparison between variable pluripotency factors and stable pluripotency factors.

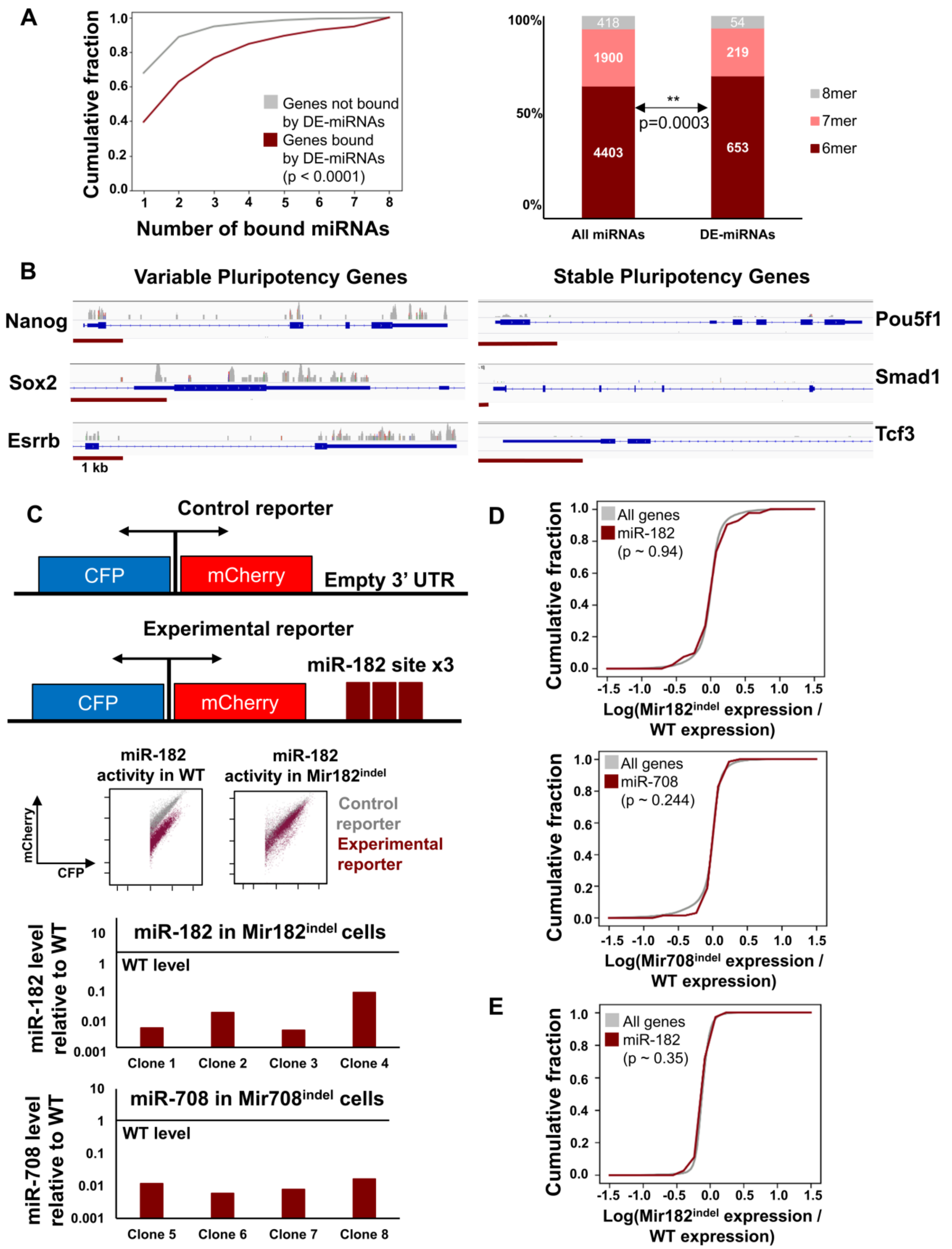
A
Cumulative fraction
Number of bound miRNAs
Genes not bound by DE-miRNAs
Genes bound by DE-miRNAs
(p < 0.0001)
100%
50%
0%
418
1900
4403
54
219
653
**
p=0.0003
8mer
7mer
6mer
All miRNAs
DE-miRNAs
B
Variable Pluripotency Genes
Stable Pluripotency Genes
Nanog
Sox2
Esrrb
1 kb
Pou5f1
Smad1
Tcf3
C
Control reporter
CFP
mCherry
Empty 3' UTR
Experimental reporter
miR-182 site x3
miR-182 activity in WT
miR-182 activity in Mir182indel
mCherry
CFP
Control reporter
Experimental reporter
miR-182 level relative to WT
miR-182 in Mir182indel cells
WT level
Clone 1
Clone 2
Clone 3
Clone 4
miR-708 level relative to WT
miR-708 in Mir708indel cells
WT level
Clone 5
Clone 6
Clone 7
Clone 8
D
Cumulative fraction
All genes
miR-182
(p ~ 0.94)
Log(Mir182indel expression / WT expression)
All genes
miR-708
(p ~ 0.244)
Log(Mir708indel expression / WT expression)
E
All genes
miR-182
(p ~ 0.35)
Log(Mir182indel expression / WT expression)

**Supplemental Figure 4: Variable microRNA binding and activity; related to Figure 3**

**A**. **Left**: CDF of number of Ago2-miRNAs binding sites for genes where at least one site is assigned to a variable DE-miRNA (red) and genes bound by Ago2-miRNA where no sites are assigned to DE-miRNAs (gray). Kolmogorov-Smirnov (K-S) p-value is shown. **Right**: Distribution of target site affinities (6mer, 7mer, or 8mer matches for the assigned miRNA seed within the cluster of Ago2-binding) for all miRNAs or variable DE-miRNAs. Hypergeometric p-value for enrichment is shown.

**B.** Ago2 iCLIP coverage and gene structure are shown for variable pluripotency genes *Nanog, Sox2, Esrrb* and less variable pluripotency genes *Pou5f1, Smad1,* and *Tcf3*. Ago2 coverage is scaled identically across all six displayed loci (linear scale). For each gene, the 3'UTR is shown along with much of the gene body. Red scale bars indicate 1 kb for each gene. Data from (Bosson et al., 2014).

**C**. **Top**: Bidirectional reporter assay for miR-182 activity in WT and *Mir182*$^{indel}$ cells. In brief, mCherry is transcribed tightly coupled to Cerulean (CFP) from a bidirectional doxycycline inducible plasmid. Three miR-182 target sites are cloned into the 3'UTR of *mCherry*. In this way, CFP levels provide a measure of total transfection and induction efficiency and mCherry levels relative to CFP provide a measure of miRNA regulation. See also (Mukherji et al., 2011; Schmiedel et al., 2015) for further details about this method. **Bottom**: miR-182 and miR-708 levels in *Mir182*$^{indel}$ cells and *Mir708*$^{indel}$ cells by quantitative PCR (using the ddCt method relative to WT ESC) respectively. Insertion/deletions were verified by sequencing across the hairpin region of the miRNA gene from amplified genomic DNA. Note that clones 3-4 & 7-8 were labeled with fluorophores at *Nanog* and *Sox2* loci, and clones 3 & 7 are those used in Fig. 5 and S6. Clones 1-2 and 5-6 were used in biological replicates for bulk RNA-sequencing of miR-182 and miR-708 targets in *Mir*$^{indel}$ cells (Fig. S4D below). Clone 2 is used for reporter activity assay shown in this figure.

**D. Top:** Expression of all genes (gray) or genes targeted by miR-182 in WT vs. *Mir182*$^{indel}$ ESC displayed as cumulative distribution function (CDF) over the logarithm of expression ratio between cell types. Three biological replicates for each condition were used in bulk RNA-sequencing (Methods). **Bottom:** Analogous ratio is displayed for all genes or those targeted by miR-708 in WT vs *Mir708*$^{indel}$ ESC. Kolmogorov-Smirnov (K-S) p-values are shown (see methods).

**E.** Expression of all genes and miR-182 target genes on average across single cells in *Mir182*$^{indel}$ vs. WT ESC. Data are displayed as CDF of logarithm of expression ratio (average *Mir182*$^{indel}$ expression / average WT expression across single cells). K-S p value is shown.

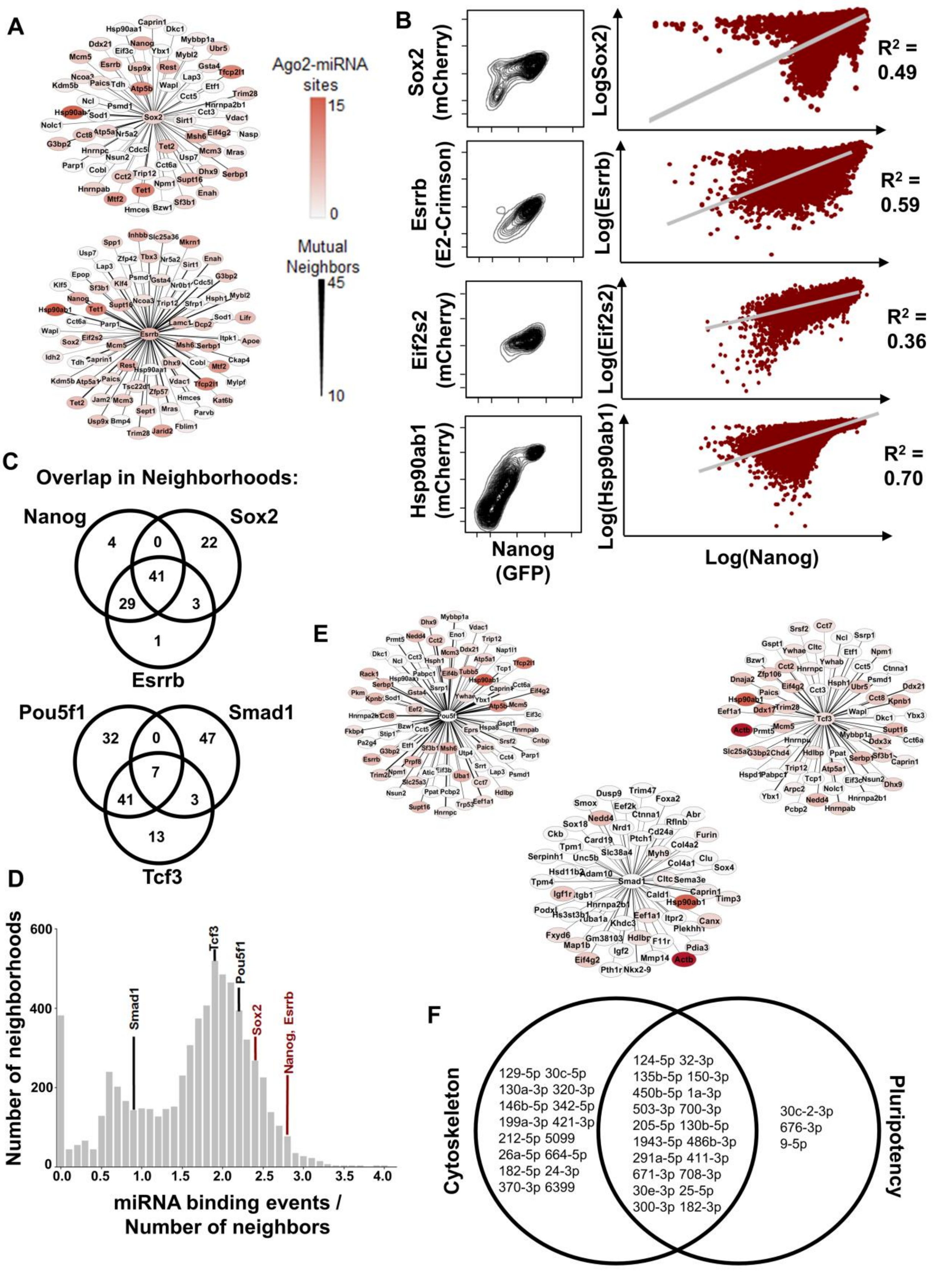
A
Ago2-miRNA sites
15
0
Mutual Neighbors
45
10
B
Sox2 (mCherry)
LogSox2
$R^2$ = 0.49
Esrrb (E2-Crimson)
Log(Esrrb)
$R^2$ = 0.59
Eif2s2 (mCherry)
Log(Eif2s2)
$R^2$ = 0.36
Hsp90ab1 (mCherry)
Log(Hsp90ab1)
$R^2$ = 0.70
Nanog (GFP)
Log(Nanog)
C
Overlap in Neighborhoods:
Nanog
Sox2
4
0
22
41
29
3
1
Esrrb
Pou5f1
Smad1
32
0
47
7
41
3
13
Tcf3
D
Number of neighborhoods
miRNA binding events / Number of neighbors
Smad1
Tcf3
Pou5f1
Sox2
Nanog, Esrrb
E
F
Cytoskeleton
Pluripotency
129-5p 30c-5p
130a-3p 320-3p
146b-5p 342-5p
199a-3p 421-3p
212-5p 5099
26a-5p 664-5p
182-5p 24-3p
370-3p 6399
124-5p 32-3p
135b-5p 150-3p
450b-5p 1a-3p
503-3p 700-3p
205-5p 130b-5p
1943-5p 486b-3p
291a-5p 411-3p
671-3p 708-3p
30e-3p 25-5p
300-3p 182-3p
30c-2-3p
676-3p
9-5p

**Supplemental Figure 5: miRNA binding to gene neighborhoods; related to Figure 4**
**A.** *Sox2* and *Esrrb* neighborhoods and degree of binding by Ago2-miRNA; analogous to Fig. 4A. Thickness of connection indicates number of mutual neighbors, and degree of red shading indicates number of Ago2-miRNA sites.
**B.** Covariation of proteins encoded by *Nanog* neighbors with Nanog. In each cell line, *Nanog* is tagged at its endogenous locus with GFP. The neighbor of *Nanog* is then tagged at its endogenous locus with the indicated fluorophore. Cells are then analyzed by flow cytometry for fluorophore levels, which approximate the tagged genes. FACS distributions are shown at left; at right are the corresponding scatter plots, where each dot represents a single cell's fluorescence levels. Best fit lines and $R^2$ values are shown.
**C**. Venn diagram showing the overlap in neighbors for variable pluripotency genes and less variable pluripotency genes. Note that 41 neighbors occur in each of the *Nanog*, *Sox2*, and *Esrrb* neighborhoods indicating these form a dense clique. In contrast, less variable pluripotency gene neighborhoods show fewer mutual neighbors, though *Pou5f1* and *Tcf3* do overlap. Note of the 41 genes overlapping in the *Nanog*/*Sox2*/*Esrrb* clique only 20 of these overlap with *Pou5f1* or *Tcf3*.
**D.** Histogram for average miRNA binding per gene (miRNA binding events / number of neighbors) for the 6,577 non-empty neighborhoods produced for all the genes present in WT scRNA-seq data. Values for *Nanog*, *Sox2*, and *Esrrb* neighborhoods are indicated in red; those for *Pou5f1*, *Smad1*, and *Tcf3* neighborhoods are shown in black.
**E.** *Pou5f1*, *Smad1*, and *Tcf3* neighborhoods and degree of binding by Ago2-miRNA; analogous to Figs. 4A and S4A.
**F.** miRNAs enriched for binding 192 cytoskeleton neighborhoods and their overlap with miRNAs enriched for binding pluripotency gene neighborhoods from Fig. 4B.

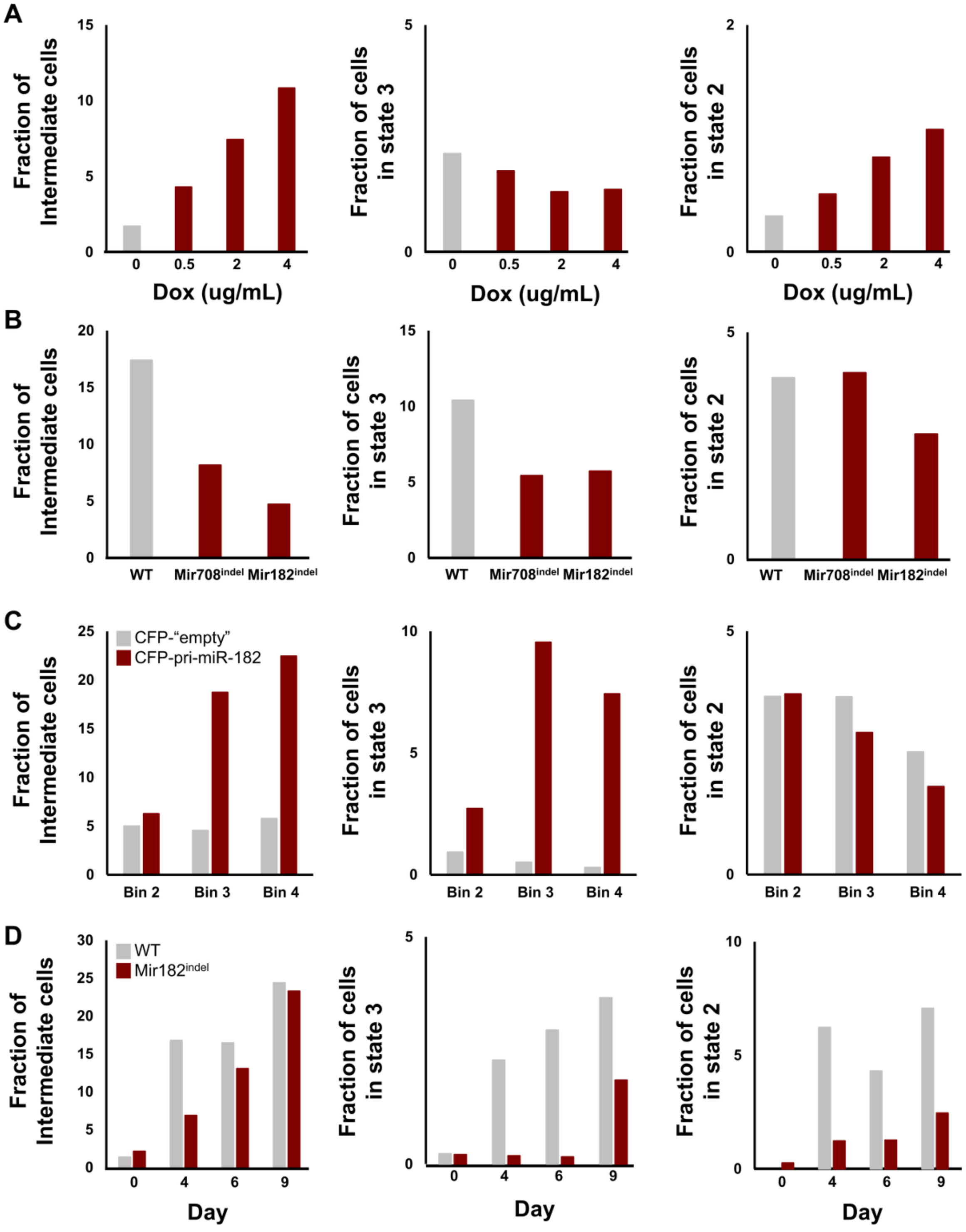
A
Fraction of Intermediate cells
Fraction of cells in state 3
Fraction of cells in state 2
Dox (ug/mL)
0
0.5
2
4
B
WT
Mir708indel
Mir182indel
C
CFP-"empty"
CFP-pri-miR-182
Bin 2
Bin 3
Bin 4
D
WT
Mir182indel
Day
0
4
6
9

**Supplemental Figure 6: Distributions amongst cell states for variable miRNA reconstituted cells; related to Figure 5**

**A**. Fraction of cells in State 2, State 3, and intermediate between States 1-3 (outside of state gating) for Ago2-inducible ESC. All cells were cultured at 1 μg/mL doxycycline continuously and then switched into the indicated amounts of doxycycline for 48 hours before measuring cell state distributions by flow cytometry. Compare to Fig. 5B.

**B**. Fraction of cells in State 2, State 3, and intermediate between States 1-3 for $\mathit{Mir}^{\mathit{indel}}$ ESC compared to WT ESC. Compare to Fig. 5C.

**C**. Fraction of cells in State 2, State 3, and intermediate between States 1-3 for $\mathit{Mir182}^{\mathit{indel}}$ cells transfected with pri-miR-182 expressing plasmid or transfected with empty control plasmid. Compare to Fig. 5E.

**D**. Fraction of cells in State 2, State 3, and intermediate between States 1-3 for WT and $\mathit{Mir182}^{\mathit{indel}}$ ESC sorted for State 1 and cultured for the indicated number of days. Compare to Fig. 5F.

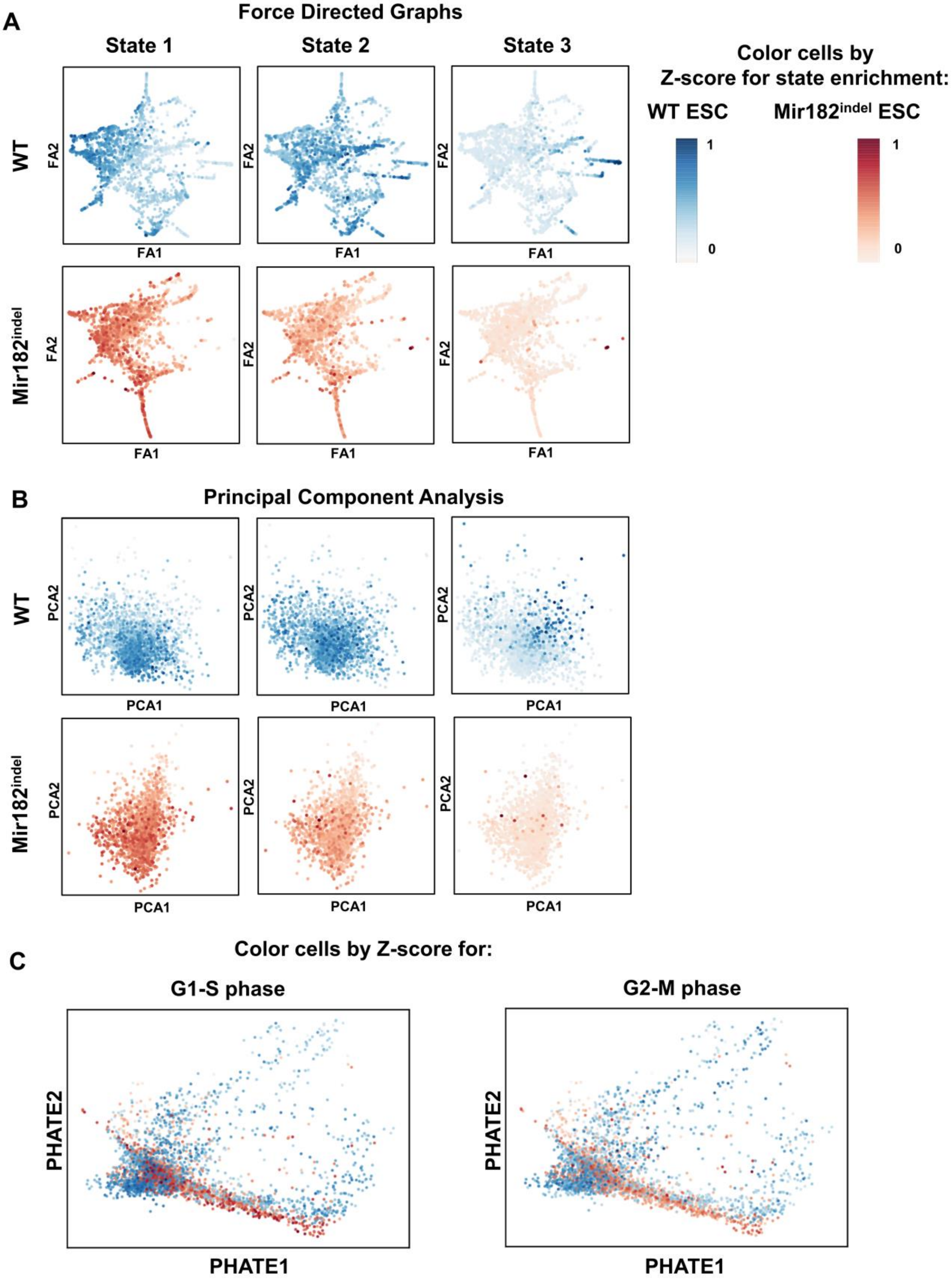
A
Force Directed Graphs
State 1
State 2
State 3
WT
Mir182indel
FA1
FA2
Color cells by
Z-score for state enrichment:
WT ESC
Mir182indel ESC
1
0
B
Principal Component Analysis
PCA1
PCA2
C
Color cells by Z-score for:
G1-S phase
G2-M phase
PHATE1
PHATE2

**Supplemental Figure 7: States defined by Nanog and Sox2 (and not stages of the cell cycle) capture a large portion of ESC state diversity; related to Figure 6**

**A.** Force directed method (Paul et al., 2015) of dimensionality reduction. WT and *Mir182*$^{indel}$ ESC were analyzed together. **Top row**: WT cells pseudocolored by Z-score for State Enrichment; **bottom row**: *Mir182*$^{indel}$ cells.

**B.** Principal Component Analysis (PCA) of WT and *Mir182*$^{indel}$ ESC is shown, plotted analogously to S7A and Fig. 6.

**C.** PHATE graphs of WT and *Mir182*$^{indel}$ ESC. Cells are placed exactly as in Fig. 6 but are pseudocolored for enrichment of expression of G1-S or G2-M specific cell cycle genes (Patel et al., 2014) instead of pseudocoloring by State 1-3 expression programs. Note the lack of definitive clustering according to either phase of the ESC cell cycle.

**References (copied from Main File)**:

Beauparlant CJ, L.F., Samb R, Lippens C, Deschenes AL, Droit A (2019). metagene: A package to produce metagene plots. R package version 2160.
Bhang, H.E., Ruddy, D.A., Krishnamurthy Radhakrishna, V., Caushi, J.X., Zhao, R., Hims, M.M., Singh, A.P., Kao, I., Rakiec, D., Shaw, P.*, et al.* (2015). Studying clonal dynamics in response to cancer therapy using high-complexity barcoding. Nat Med *21*, 440-448.
Boroviak, T., Loos, R., Lombard, P., Okahara, J., Behr, R., Sasaki, E., Nichols, J., Smith, A., and Bertone, P. (2015). Lineage-Specific Profiling Delineates the Emergence and Progression of Naive Pluripotency in Mammalian Embryogenesis. Dev Cell *35*, 366-382.
Bosia, C., Pagnani, A., and Zecchina, R. (2013). Modelling Competing Endogenous RNA Networks. PLoS One *8*, e66609.
Bosson, A.D., Zamudio, J.R., and Sharp, P.A. (2014). Endogenous miRNA and target concentrations determine susceptibility to potential ceRNA competition. Mol Cell *56*, 347-359.
Chambers, I., Silva, J., Colby, D., Nichols, J., Nijmeijer, B., Robertson, M., Vrana, J., Jones, K., Grotewold, L., and Smith, A. (2007). Nanog safeguards pluripotency and mediates germline development. Nature *450*, 1230-1234.
Chang, H.H., Hemberg, M., Barahona, M., Ingber, D.E., and Huang, S. (2008). Transcriptome-wide noise controls lineage choice in mammalian progenitor cells. Nature *453*, 544-547.
Chen, Q., Shi, J., Tao, Y., and Zernicka-Goetz, M. (2018). Tracing the origin of heterogeneity and symmetry breaking in the early mammalian embryo. Nat Commun *9*, 1819.
Chen, S., Sanjana, N.E., Zheng, K., Shalem, O., Lee, K., Shi, X., Scott, D.A., Song, J., Pan, J.Q., Weissleder, R.*, et al.* (2015). Genome-wide CRISPR screen in a mouse model of tumor growth and metastasis. Cell *160*, 1246-1260.
Cho, W.K., Spille, J.H., Hecht, M., Lee, C., Li, C., Grube, V., and Cisse, II (2018). Mediator and RNA polymerase II clusters associate in transcription-dependent condensates. Science *361*, 412-415.
Chong, S., Dugast-Darzacq, C., Liu, Z., Dong, P., Dailey, G.M., Cattoglio, C., Heckert, A., Banala, S., Lavis, L., Darzacq, X.*, et al.* (2018). Imaging dynamic and selective low-complexity domain interactions that control gene transcription. Science *361*.
Del Giudice, M., Bo, S., Grigolon, S., and Bosia, C. (2018). On the role of extrinsic noise in microRNA-mediated bimodal gene expression. PLoS Comput Biol *14*, e1006063.
Deng, Q., Ramskold, D., Reinius, B., and Sandberg, R. (2014). Single-cell RNA-seq reveals dynamic, random monoallelic gene expression in mammalian cells. Science *343*, 193-196.
Dowen, J.M., Fan, Z.P., Hnisz, D., Ren, G., Abraham, B.J., Zhang, L.N., Weintraub, A.S., Schujiers, J., Lee, T.I., Zhao, K.*, et al.* (2014). Control of cell identity genes occurs in insulated neighborhoods in mammalian chromosomes. Cell *159*, 374-387.
Eldar, A., and Elowitz, M.B. (2010). Functional roles for noise in genetic circuits. Nature *467*, 167-173.
Fukaya, T., Lim, B., and Levine, M. (2016). Enhancer Control of Transcriptional Bursting. Cell *166*, 358-368.
Gardner, R.L. (2001). Specification of embryonic axes begins before cleavage in normal mouse development. Development *128*, 839-847.
Garg, S., and Sharp, P.A. (2016). GENE EXPRESSION. Single-cell variability guided by microRNAs. Science *352*, 1390-1391.
Gierahn, T.M., Wadsworth, M.H., 2nd, Hughes, T.K., Bryson, B.D., Butler, A., Satija, R., Fortune, S., Love, J.C., and Shalek, A.K. (2017). Seq-Well: portable, low-cost RNA sequencing of single cells at high throughput. Nat Methods *14*, 395-398.

Gillespie, D.T. (1976). General Method for Numerically Simulating Stochastic Time Evolution of Coupled Chemical-Reactions. J Comput Phys *22*, 403-434.
Goolam, M., Scialdone, A., Graham, S.J.L., Macaulay, I.C., Jedrusik, A., Hupalowska, A., Voet, T., Marioni, J.C., and Zernicka-Goetz, M. (2016). Heterogeneity in Oct4 and Sox2 Targets Biases Cell Fate in 4-Cell Mouse Embryos. Cell *165*, 61-74.
Gu, Z., Eils, R., and Schlesner, M. (2016). Complex heatmaps reveal patterns and correlations in multidimensional genomic data. Bioinformatics *32*, 2847-2849.
Harrison, S.E., Sozen, B., Christodoulou, N., Kyprianou, C., and Zernicka-Goetz, M. (2017). Assembly of embryonic and extraembryonic stem cells to mimic embryogenesis in vitro. Science *356*.
Hnisz, D., Abraham, B.J., Lee, T.I., Lau, A., Saint-Andre, V., Sigova, A.A., Hoke, H.A., and Young, R.A. (2013). Super-enhancers in the control of cell identity and disease. Cell *155*, 934-947.
Hnisz, D., Shrinivas, K., Young, R.A., Chakraborty, A.K., and Sharp, P.A. (2017). A Phase Separation Model for Transcriptional Control. Cell *169*, 13-23.
Ju, Y.S., Martincorena, I., Gerstung, M., Petljak, M., Alexandrov, L.B., Rahbari, R., Wedge, D.C., Davies, H.R., Ramakrishna, M., Fullam, A.*, et al.* (2017). Somatic mutations reveal asymmetric cellular dynamics in the early human embryo. Nature *543*, 714-718.
Kagey, M.H., Newman, J.J., Bilodeau, S., Zhan, Y., Orlando, D.A., van Berkum, N.L., Ebmeier, C.C., Goossens, J., Rahl, P.B., Levine, S.S.*, et al.* (2010). Mediator and cohesin connect gene expression and chromatin architecture. Nature *467*, 430-435.
Klein, A.M., Mazutis, L., Akartuna, I., Tallapragada, N., Veres, A., Li, V., Peshkin, L., Weitz, D.A., and Kirschner, M.W. (2015). Droplet barcoding for single-cell transcriptomics applied to embryonic stem cells. Cell *161*, 1187-1201.
Kumar, R.M., Cahan, P., Shalek, A.K., Satija, R., DaleyKeyser, A., Li, H., Zhang, J., Pardee, K., Gennert, D., Trombetta, J.J.*, et al.* (2014). Deconstructing transcriptional heterogeneity in pluripotent stem cells. Nature *516*, 56-61.
Kurotaki, Y., Hatta, K., Nakao, K., Nabeshima, Y., and Fujimori, T. (2007). Blastocyst axis is specified independently of early cell lineage but aligns with the ZP shape. Science *316*, 719-723.
Larsson, A.J.M., Johnsson, P., Hagemann-Jensen, M., Hartmanis, L., Faridani, O.R., Reinius, B., Segerstolpe, A., Rivera, C.M., Ren, B., and Sandberg, R. (2019). Genomic encoding of transcriptional burst kinetics. Nature *565*, 251-254.
Levine, M., and Tjian, R. (2003). Transcription regulation and animal diversity. Nature *424*, 147-151.
Li, A., and Horvath, S. (2007). Network neighborhood analysis with the multi-node topological overlap measure. Bioinformatics *23*, 222-231.
Marson, A., Levine, S.S., Cole, M.F., Frampton, G.M., Brambrink, T., Johnstone, S., Guenther, M.G., Johnston, W.K., Wernig, M., Newman, J.*, et al.* (2008). Connecting microRNA genes to the core transcriptional regulatory circuitry of embryonic stem cells. Cell *134*, 521-533.
Martirosyan, A., Figliuzzi, M., Marinari, E., and De Martino, A. (2016). Probing the Limits to MicroRNA-Mediated Control of Gene Expression. PLoS Comput Biol *12*, e1004715.
Medeiros, L.A., Dennis, L.M., Gill, M.E., Houbaviy, H., Markoulaki, S., Fu, D., White, A.C., Kirak, O., Sharp, P.A., Page, D.C.*, et al.* (2011). Mir-290-295 deficiency in mice results in partially penetrant embryonic lethality and germ cell defects. Proc Natl Acad Sci U S A *108*, 14163-14168.
Melton, C., Judson, R.L., and Blelloch, R. (2010). Opposing microRNA families regulate self-renewal in mouse embryonic stem cells. Nature *463*, 621-626.
Moon, K.R., van Dijk, D., Wang, Z., Gigante, S., Burkhardt, D.B., Chen, W.S., Yim, K., van den Elzen, A., Hirn, M.J., Coifman, R.R.*, et al.* (2019). Visualizing Structure and Transitions for Biological Data Exploration. bioRxiv.
Motosugi, N., Bauer, T., Polanski, Z., Solter, D., and Hiiragi, T. (2005). Polarity of the mouse embryo is established at blastocyst and is not prepatterned. Genes Dev *19*, 1081-1092.

Mukherji, S., Ebert, M.S., Zheng, G.X., Tsang, J.S., Sharp, P.A., and van Oudenaarden, A. (2011). MicroRNAs can generate thresholds in target gene expression. Nat Genet *43*, 854-859.
Noorbakhsh, J., Lang, A.H., and Mehta, P. (2013). Intrinsic noise of microRNA-regulated genes and the ceRNA hypothesis. PLoS One *8*, e72676.
Patel, A.P., Tirosh, I., Trombetta, J.J., Shalek, A.K., Gillespie, S.M., Wakimoto, H., Cahill, D.P., Nahed, B.V., Curry, W.T., Martuza, R.L.*, et al.* (2014). Single-cell RNA-seq highlights intratumoral heterogeneity in primary glioblastoma. Science *344*, 1396-1401.
Paul, F., Arkin, Y., Giladi, A., Jaitin, D.A., Kenigsberg, E., Keren-Shaul, H., Winter, D., Lara-Astiaso, D., Gury, M., Weiner, A.*, et al.* (2015). Transcriptional Heterogeneity and Lineage Commitment in Myeloid Progenitors. Cell *163*, 1663-1677.
Piotrowska-Nitsche, K., Perea-Gomez, A., Haraguchi, S., and Zernicka-Goetz, M. (2005). Four-cell stage mouse blastomeres have different developmental properties. Development *132*, 479-490.
Raj, A., and van Oudenaarden, A. (2008). Nature, nurture, or chance: stochastic gene expression and its consequences. Cell *135*, 216-226.
Rivron, N.C., Frias-Aldeguer, J., Vrij, E.J., Boisset, J.C., Korving, J., Vivie, J., Truckenmuller, R.K., van Oudenaarden, A., van Blitterswijk, C.A., and Geijsen, N. (2018). Blastocyst-like structures generated solely from stem cells. Nature *557*, 106-111.
Sabari, B.R., Dall'Agnese, A., Boija, A., Klein, I.A., Coffey, E.L., Shrinivas, K., Abraham, B.J., Hannett, N.M., Zamudio, A.V., Manteiga, J.C.*, et al.* (2018). Coactivator condensation at super-enhancers links phase separation and gene control. Science *361*.
Schmiedel, J.M., Klemm, S.L., Zheng, Y., Sahay, A., Bluthgen, N., Marks, D.S., and van Oudenaarden, A. (2015). Gene expression. MicroRNA control of protein expression noise. Science *348*, 128-132.
Shahbazi, M.N., Scialdone, A., Skorupska, N., Weberling, A., Recher, G., Zhu, M., Jedrusik, A., Devito, L.G., Noli, L., Macaulay, I.C.*, et al.* (2017). Pluripotent state transitions coordinate morphogenesis in mouse and human embryos. Nature *552*, 239-243.
Singer, Z.S., Yong, J., Tischler, J., Hackett, J.A., Altinok, A., Surani, M.A., Cai, L., and Elowitz, M.B. (2014). Dynamic heterogeneity and DNA methylation in embryonic stem cells. Mol Cell *55*, 319-331.
Singh, A., Razooky, B., Cox, C.D., Simpson, M.L., and Weinberger, L.S. (2010). Transcriptional bursting from the HIV-1 promoter is a significant source of stochastic noise in HIV-1 gene expression. Biophys J *98*, L32-34.
Solter, D. (2016). Preformation Versus Epigenesis in Early Mammalian Development. Curr Top Dev Biol *117*, 377-391.
Stelzer, Y., Shivalila, C.S., Soldner, F., Markoulaki, S., and Jaenisch, R. (2015). Tracing dynamic changes of DNA methylation at single-cell resolution. Cell *163*, 218-229.
Stewart-Ornstein, J., and Lahav, G. (2016). Dynamics of CDKN1A in Single Cells Defined by an Endogenous Fluorescent Tagging Toolkit. Cell Rep *14*, 1800-1811.
Su, H., Trombly, M.I., Chen, J., and Wang, X. (2009). Essential and overlapping functions for mammalian Argonautes in microRNA silencing. Genes Dev *23*, 304-317.
Suzuki, H.I., Young, R.A., and Sharp, P.A. (2017). Super-Enhancer-Mediated RNA Processing Revealed by Integrative MicroRNA Network Analysis. Cell *168*, 1000-1014 e1015.
Torres-Padilla, M.E., Parfitt, D.E., Kouzarides, T., and Zernicka-Goetz, M. (2007). Histone arginine methylation regulates pluripotency in the early mouse embryo. Nature *445*, 214-218.
Wang, Y., Medvid, R., Melton, C., Jaenisch, R., and Blelloch, R. (2007). DGCR8 is essential for microRNA biogenesis and silencing of embryonic stem cell self-renewal. Nat Genet *39*, 380-385.
White, M.D., Angiolini, J.F., Alvarez, Y.D., Kaur, G., Zhao, Z.W., Mocskos, E., Bruno, L., Bissiere, S., Levi, V., and Plachta, N. (2016). Long-Lived Binding of Sox2 to DNA Predicts Cell Fate in the Four-Cell Mouse Embryo. Cell *165*, 75-87.

Whyte, W.A., Orlando, D.A., Hnisz, D., Abraham, B.J., Lin, C.Y., Kagey, M.H., Rahl, P.B., Lee, T.I., and Young, R.A. (2013). Master transcription factors and mediator establish super-enhancers at key cell identity genes. Cell *153*, 307-319.

Wolf, F.A., Angerer, P., and Theis, F.J. (2018). SCANPY: large-scale single-cell gene expression data analysis. Genome Biol *19*, 15.

Zamudio, J.R., Kelly, T.J., and Sharp, P.A. (2014). Argonaute-bound small RNAs from promoter-proximal RNA polymerase II. Cell *156*, 920-934.
